# Molecular Regulation of Invasive Protrusion Formation at the Mammalian Fusogenic Synapse

**DOI:** 10.1101/2023.11.27.568897

**Authors:** Yue Lu, Tezin Walji, Benjamin Ravaux, Pratima Pandey, Bing Li, Kevin H. Lam, Ruihui Zhang, David J. Goldhamer, Rong Li, David W. Schmidtke, Duojia Pan, Elizabeth H. Chen

## Abstract

Invasive membrane protrusions play a central role in a variety of cellular processes. Unlike filopodia, invasive protrusions are mechanically stiff and propelled by branched actin polymerization. However, how branched actin filaments are organized to create finger-like invasive protrusions remains a longstanding question in cell biology. Here, by examining the mammalian fusogenic synapse, where invasive protrusions are generated to promote cell membrane juxtaposition and fusion, we have uncovered the mechanism underlying invasive protrusion formation. We show that two Arp2/3 nucleation promoting factors (NPFs), WAVE and N-WASP, exhibit distinct and complementary localization patterns in the protrusions. While WAVE is at the leading edge, N-WASP is recruited by its interacting protein, WIP, to the shaft of the protrusion. During protrusion growth, new branched actin filaments are polymerized at the periphery of the shaft and crosslinked to preexisting actin bundles by the “pioneer” actin-bundling protein dynamin. The thickened actin bundles are further stabilized by WIP, which functions as a WH2 domain-mediated actin-bundling protein. Disrupting any of these components results in defective protrusions and myoblast fusion in cultured cells and/or in mouse embryos. Thus, our study has revealed the intricate spatiotemporal coordination between two NPFs and two actin-bundling proteins in creating invasive protrusions and has general implications in understanding protrusion formation in many cellular processes beyond cell-cell fusion.

## Introduction

Actin-based invasive membrane protrusions are essential for many cellular processes, including cancer metastasis, leukocyte diapedesis or transendothelial migration, immunological synapse maturation, uterine-vulval attachment, and cell-cell fusion^1-5^. In cancer metastasis, actin-rich structures called invadopodia are required for cancer cell extravasation by invading and degrading the basement membrane^6-8^. Leukocytes utilize actin-rich podosomes to invade endothelia cells during their transendothelial migration^2,8^. At the immunological synapses, cytotoxic T lymphocytes also generate actin-rich protrusions to deform the target cell surface and enhance cellular toxicity^3^. During *C. elegans* development, a specialized uterine cell, anchor cell, uses invadopodia to breach the basement membrane and initiate uterine-vulval attachment^4^.

Cell-cell fusion is required for the development and regeneration of multicellular organisms^9-11^. Major insights into the mechanisms of cell-cell fusion came from studies of skeletal muscle cell fusion, in which mononucleated myoblasts fuse to form multinucleated contractile muscle fibers^5,12-14^. Work in *Drosophila* myoblast fusion led to the discovery of the asymmetric fusogenic synapse, where a cell generates an F-actin-enriched podosome-like structure (PLS) that projects invasive membrane protrusions into its fusion partner to promote cell membrane juxtaposition and fusion^13-16^. Recent studies in zebrafish embryos have demonstrated that fusion between fast muscle cells is mediated by similar F-actin-propelled invasive protrusions^17^. Besides myoblast fusion, induced fusion between cultured *Drosophila* cells of hemocyte origin also depends on invasive protrusions to bring cell membranes into close proximity for fusogen engagement and membrane merger. Thus, both muscle and non-muscle cells use invasive protrusions as a general mechanism to facilitate cell membrane fusion. The essential function of invasive protrusions *in vivo* has been highlighted by *Drosophila* myoblast fusion mutants in which the invasiveness of the protrusions is compromised^15,19-21^.

In *Drosophila*, invasive protrusions at the fusogenic synapse are propelled by Arp2/3- mediated branched actin polymerization^13^. Two actin nucleation-promoting factors (NPFs) for the Arp2/3 complex, WASP and WAVE (also known as Scar)^22^, have redundant functions in activating actin polymerization at the fusogenic synapse^15^. WASP and WASP-interacting protein (WIP, also known as Solitary or Sltr) colocalize with the F-actin core of the PLS^15,19,23^, whereas the localization of WAVE remains unclear. Genetic analyses in *Drosophila* showed that the WASP/WIP complex, but not the pentameric WAVE complex, is required for the invasiveness of the protrusions^15,19^. How WASP and WAVE, which have similar biochemical functions towards activating the Arp2/3 complex, exert distinct functions in regulating the invasiveness of protrusions is unclear.

To generate invasive protrusions of ∼250 nm diameter^15^, the branched actin filaments must be organized into mechanically stiff actin bundles. A recent study revealed an essential function for the large GTPase dynamin as a multi-filament actin-bundling protein in *Drosophila* myoblast fusion^21^. Dynamin is enriched within the PLS at the fusogenic synapse and promotes the invasiveness of membrane protrusions by bundling actin filaments while forming a helical structure^21^. At its full capacity, a single *Drosophila* dynamin helix can bundle 12 actin filaments on the outer rim of the helix, whereas its mammalian counterpart can bundle 16 actin filaments, making dynamin a highly efficient, unique actin-bundling protein^21^. Once a dynamin helix is fully assembled, it undergoes rapid disassembly induced by GTP hydrolysis, leaving behind loosely aligned actin filaments and releasing dynamin dimers back into the cytosol for new rounds of actin bundling^21^. Despite the genetic and biochemical analyses, how dynamin spatially and temporally bundles actin filaments during invasive protrusion formation remains unclear. Furthermore, since dynamin helices disassemble after bundling actin filaments, how the loose actin bundles are stabilized is completely unknown.

Compared to *Drosophila* and zebrafish myoblast fusion, the molecular and cellular events mediating mammalian myoblast fusion are poorly understood. To date, several actin cytoskeletal regulators have been implicated in mammalian myoblast fusion by loss-of-function analyses in mouse embryos, such as N-WASP and its activator Cdc42, an activator for the WAVE complex (Rac1), and the bi-partite guanine nucleotide exchange factor for Rac1 (Dock180/Dock1 and Elmo)^24-27^, or in C2C12 cells, such as a subunit of the WAVE complex (Nap1)^28^. In addition, two muscle-specific transmembrane (TM) fusogenic proteins, myomaker (MymK) and myomixer (MymX)/myomerger/minion, are required for myoblast fusion *in vivo* and can induce cell-cell fusion when co-expressed in fibroblasts^29-32^. Despite their fusogenic activities, MymK and MymX require a functional actin cytoskeleton to induce cell-cell fusion^29,32^, highlighting the importance of the actin cytoskeleton in the fusion process. However, despite the identification of actin regulators in mammalian myoblast fusion and the observed actin enrichment and/or membrane protrusions prior to fusion in cultured mammalian muscle cells^33-36^, the localization of the actin regulators relative to the fusion sites and how these actin regulators coordinate with one another to build the fusion machinery have yet to be revealed.

In this study, we demonstrate that mammalian myoblasts fusion is mediated by F-actin-propelled invasive membrane protrusions. Formation of the invasive protrusions is orchestrated by multiple branched actin polymerization regulators and actin bundling proteins. While WAVE2 is localized at the leading edge of the invasive structure, N-WASP is recruited by its interacting protein, WIP2, to the shaft of the protrusion. During protrusion growth, new branched actin filaments are polymerized at the periphery of the shaft and crosslinked to preexisting actin bundles by dynamin. The thickened actin bundles are then further stabilized by WIP2, which functions as a WH2 domain-mediated actin-bundling protein. Taken together, our study has revealed the intricate spatiotemporal coordination between two NPFs and two actin-bundling proteins in creating invasive protrusions at the mammalian fusogenic synapse, and shed light on the general principles of how mechanically stiff protrusions form in various cellular contexts beyond cell-cell fusion.

## Results

### Upregulation of actin polymerization prior to myoblast fusion

In order to assess the potential role for actin cytoskeleton in mammalian myoblast fusion, we examined actin polymerization in muscle cells prior to fusion. Phalloidin staining of a mouse myoblast cell line, C2C12 cells, at different time points after switching from growth media (GM; 10% fetal bovine serum) to differentiation medium (DM; 2% horse serum) revealed a gradual increase in the number of muscle cells with high F-actin content (Fig. 1a,b). The increased amount of F-actin correlated with the enhanced expression of a myogenic regulatory factor (MRF), myogenin (MyoG), and a muscle structural protein, muscle myosin heavy chain (MHC) (Fig. 1a,b), suggesting that actin polymerization correlated with muscle cell differentiation. Furthermore, overexpressing MyoG in a non-myogenic cell line, the human osteosarcoma U2OS cells, significantly increased the cellular F-actin content (Extended Data Fig. 1a), demonstrating that actin polymerization is activated by MyoG. In order to examine the potential link between actin polymerization and myoblast fusion, which is an indispensable step in muscle cell differentiation, we seeded the C2C12 cells on micropatterns (250 μm in diameter) to restrict a small number of cells within a microscopic field and followed them during the differentiation process using live cell imaging (Fig. 1c,d). After two days in DM, we used SiR-Actin, a cell permeable and specific probe for F-actin^37^, to label F-actin for 30 min before imaging. Strikingly, only myoblasts with high SiR-Actin labeling underwent fusion, consistent with a role for the actin cytoskeleton in the fusion process (Fig. 1e and Supplemental Video 1). Taken together, these data suggest that actin polymerization is upregulated prior to myoblast fusion during myogenic differentiation.

**Figure 1.**
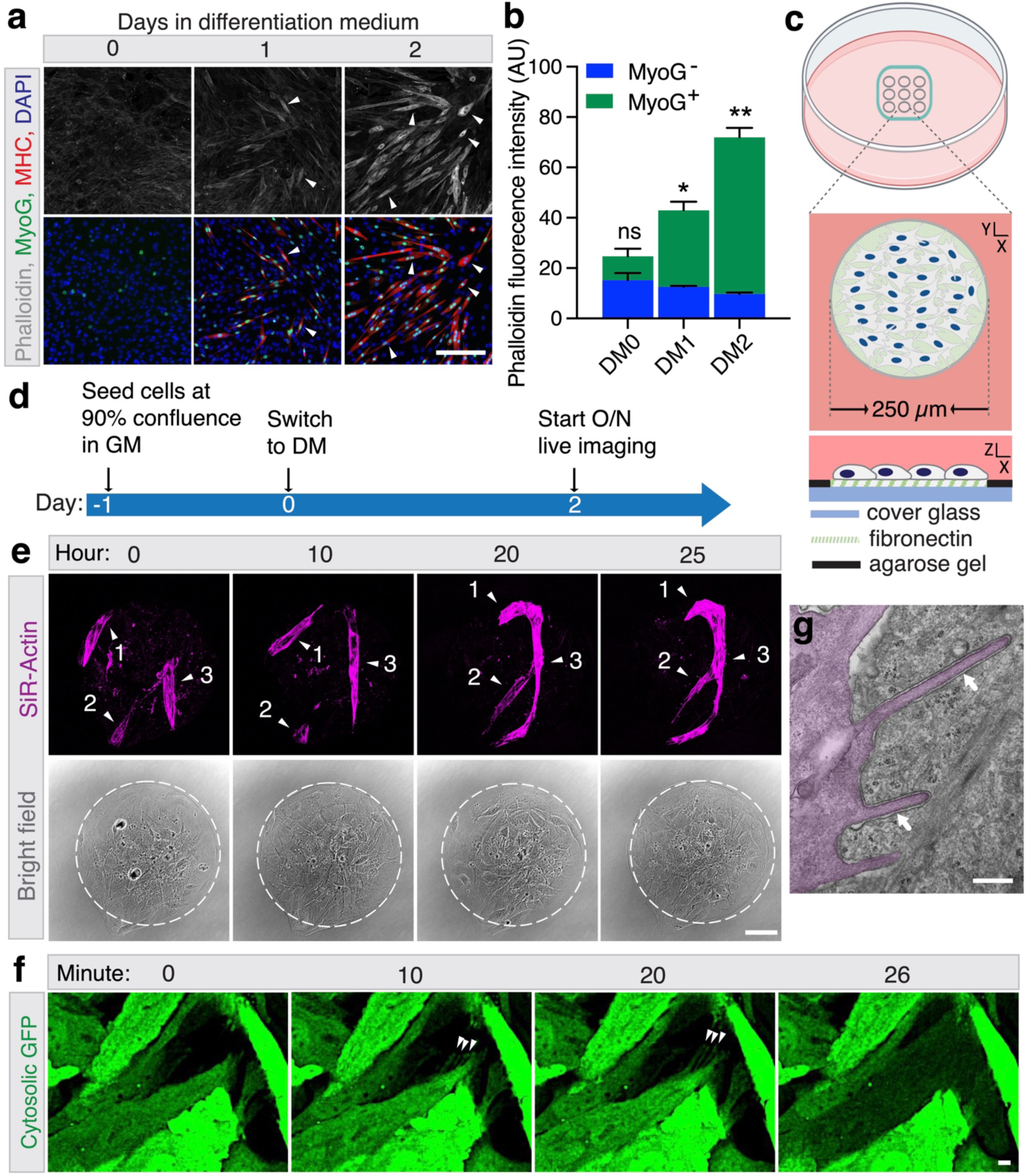
Myoblast fusion coincides with increased actin polymerization and membrane protrusion formation during mouse myoblast differentiation. **(a)** The gradual increase in the F-actin level during mouse myoblast differentiation. Wild-type C2C12 cells cultured in differentiation medium (DM) were immunostained with anti-myogenin (MyoG), anti-myosin heavy chain (MHC) and phalloidin at the indicated time points. Note the gradual increase in F-actin intensity in the cells over time. A few randomly selected F-actin^high^ cells are indicated by arrowheads. **(b)** Quantification of the phalloidin signal intensity in cells shown in (a). MyoG^-^ or MyoG^+^ cells (n ≥ 18) at each indicated time point were measured for phalloidin intensity per experiment. Three independent experiments were performed. Mean ± s.d. values are shown in the bar graph, and significance was determined by two-tailed student’s t-test. *: *p* < 0.05, **: *p* < 0.01, ns: not significant. **(c)** Schematic diagrams of the micropatterns for live cell imaging. Each micropattern is a circular fibronectin layer with a diameter of 250 μm on the glass coverslip. **(d)** Schematic diagrams of C2C12 cell culture in DM for live imaging. C2C12 cells were seeded on the micropatterns shown in (c) with growth medium (GM). Upon 100% confluency, the cells were differentiated in DM for 48 hours and subsequently subjected to confocal microscopy for live imaging. **(e)** Still images of fusion events of C2C12 cells labeled by SiR-Actin. C2C12 cells on micropatterns as describe in (d) were incubated with SiR-Actin in DM for 30 minutes to label F-actin in the cells, and immediately subjected to live imaging by confocal microscopy (see Supplemental Video 1). Three fusing myoblasts are marked as 1, 2 and 3 and indicated with arrowheads. Three independent experiments were performed with similar results. **(f)** Still images of a fusion event between GFP^+^ and GFP^-^ C2C12 cells. C2C12 cells with or without cytosolic GFP were mixed, cultured and subjected to live imaging on micropatterns as described in **(d)**. Invasive membrane protrusions at the site of fusion are indicated by arrowheads (see Supplemental Video 2). *n* = 22 fusion events were observed with similar results. **(g)** Transmission electron micrographs (TEM) of C2C12 cells cultured for 48 hours in DM. Invasive membrane protrusions at the cell-cell contact site are indicated by arrows. The invading fusion partner is pseudo-colored in magenta. *n* = 15 cell-cell contact sites were observed with similar results. Scale bars, 50 μm **(a)**, 55 μm **(e)**, 5 μm **(f)**, and 500 nm **(g)**.

### Myoblast fusion is mediated by F-actin-propelled invasive membrane protrusions

To observe the actin cytoskeletal dynamics at the sites of mammalian myoblast fusion, we performed live imaging of co-cultured C2C12 cells with or without cytosolic GFP at day two in DM. GFP^+^ finger-like membrane protrusions were observed invading into GFP^−^ fusion partners, followed by cell fusion indicated by GFP transfer (Fig. 1f and Supplemental Video 2). In addition, live imaging of fusion between F-tractin-mCherry-expressing C2C12 cells with or without cytosolic GFP revealed F-actin enrichment within the invasive structure immediately before GFP transfer between the fusing cells (Extended Data Fig. 1b and Supplemental Video 3). Consistent with this, transmission electron microscopy (TEM) analyses of differentiated C2C12 cells revealed actin-rich invasive membrane protrusions at cell-cell contact sites (Fig. 1g). Moreover, F-actin enrichment was also observed at the fusion sites of differentiated mouse satellite cells (Extended Data Fig. 1c,d and Supplemental Video 4).

Since myoblast fusion occurs in confluent C2C12 cells, each of which has high cytosolic actin content and wide cell-cell contact zones with neighboring cells, it is challenging to identify the fusion sites and visualize the fine details of the actin-enriched structures where fusion occurs. Interestingly, overexpressing an MRF, e.g. MyoD or MyoG, in sparsely localized C2C12 cells in GM could induce efficient myoblast fusion (Fig. 2a,b), allowing clearer visualization of the actin structures at the fusion sites. Live imaging of MyoD-overexpressing (MyoD^OE^) cells revealed dynamic buildup of protrusions projected from invading cells into their fusion partners (Fig. 2c-e and Supplemental Video 5). In most cases, an invading cell first generated a single protrusion (Fig. 2e, minute 4 and 6), which rapidly grew into a larger invasive structure containing multiple protrusions (Fig. 2e, minute 8 and 10). At the completion of fusion indicated by the transfer of farnesylation signal (FS)-GFP from the invading cell to the receiving cell, the F-actin enriched invasive structure rapidly dissolved (Fig. 2d, minute 12 and Fig. 2e, minute 12). Consistent with live imaging, TEM analysis of MyoD^OE^ cells also revealed multiple invasive protrusions, as in C2C12 cells (Fig. 1g), propelled by densely packed actin bundles (Fig. 2f). Taken together, these analyses demonstrate that F-actin-propelled finger-like invasive protrusions mediate mammalian myoblasts fusion and we will refer the sites of mammalian myoblast fusion as the asymmetric fusogenic synapse hereafter.

**Figure 2.**
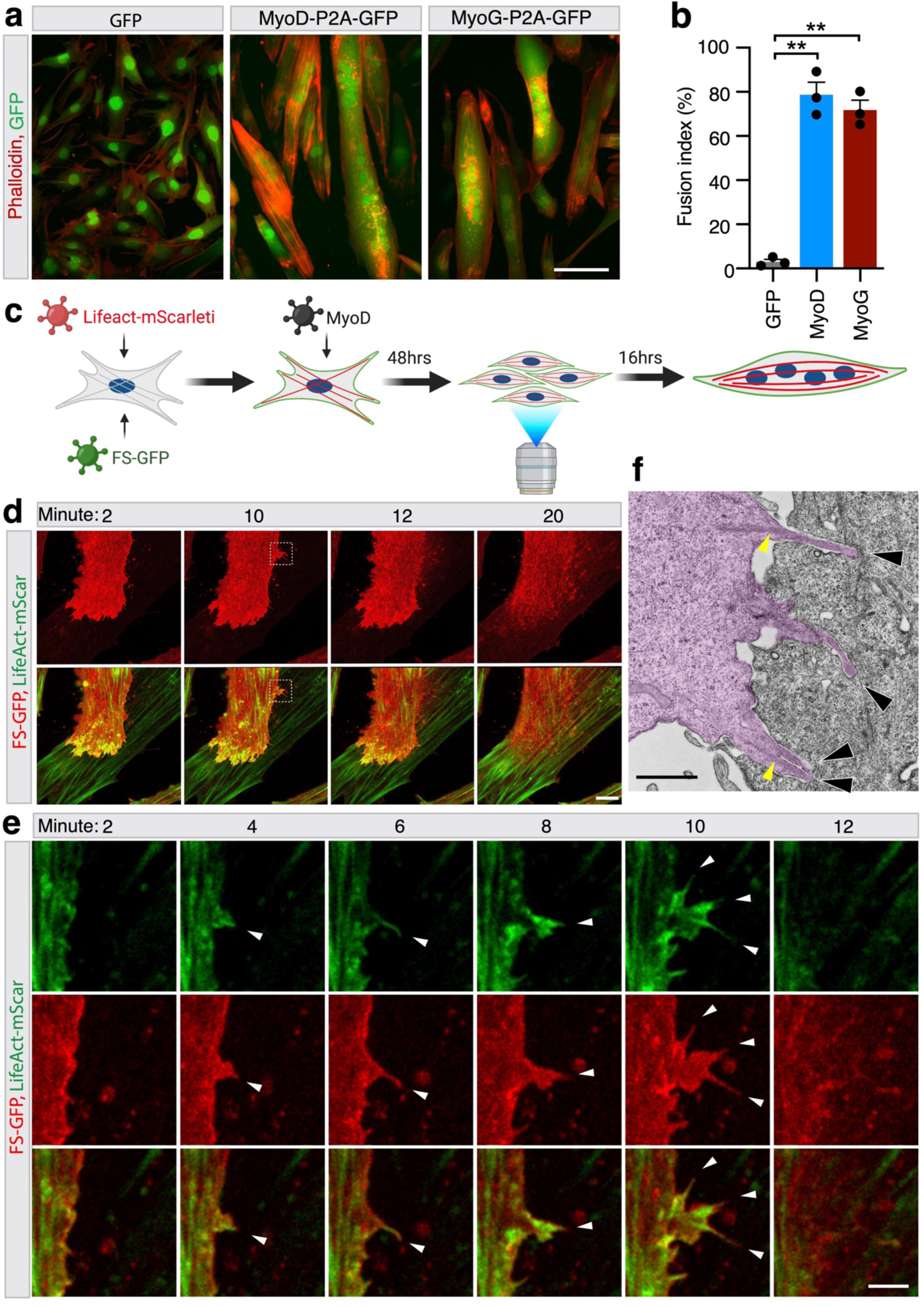
Dynamics and ultrastructure of invasive F-actin protrusions at mouse myoblast fusogenic synapse. **(a)**MyoD or MyoG overexpression (OE) induces fusion of C2C12 cells at low cell density in GM. GFP, MyoD-P2A-GFP or MyoG-P2A-GFP were transduced in C2C12 cells by retrovirus infection. 48 hours post-infection, the cells were stained with phalloidin. Note the multinucleated myofibers in MyoD^OE^ and MyoG^OE^ cells, but not in the control. **(b)** Quantification of the fusion index of the cells shown in (**a**). Fusion index was calculated as the percentage of nuclei in MHC^+^ myotubes (≥ 3 nuclei) vs. total nuclei. Three independent experiments were performed. Mean ± s.d. values are shown in the bar graph, and significance was determined by two-tailed student’s t-test. **: *p* < 0.01. **(c)** Schematic diagrams of MyoD^OE^ C2C12 cell culture in GM for live imaging. A stable C2C12 cell line co-expressing LifeAct-mScarleti (mScar) and farnesylation signal (FS)- GFP (labeling was generated by retrovirus infection) was seeded at 30% confluency in GM. After 24 hours, the cells were infected with retrovirus containing MyoD. 48 hours after MyoD overexpression, the culture medium was replaced with fresh GM and the cells were subjected to live imaging by confocal microscopy for 16 hours. **(d)** Still images of a fusion event between two MyoD^OE^ C2C12 cells. MyoD^OE^ C2C12 cells were imaged as described in (**c**). Note the transfer of FS-GFP fluorescence (12 minute) at the location of the invasive protrusions (10 minute, dotted box), indicating that a fusion pore formed at this location. Boxed area is enlarged in panel (**e**) (see Supplemental Video 5). *n* = 51 fusion events were observed with similar results. **(e)** Enlarged view of boxed area in (**d**). The finger-like protrusions are indicated by arrowheads. **(f)** TEM of C2C12 cells at 48 hours post MyoD overexpression. The invading fusion partner is pseudo-colored in magenta. Invasive protrusions are indicated by black arrowheads. Note the dense actin bundles along the shaft of the invasive protrusions (yellow arrowheads). *n* = 13 fusogenic synapses were observed with similar results. Scale bar, 100 μm (**a**), 10 μm (**d**), 3 μm (**e**), and 2 μm (**f**).

### Branched actin polymerization regulators promote mammalian myoblast fusion

To ask what type of actin nucleators are involved in actin polymerization during mammalian myoblast fusion, we pharmacologically inhibited overall actin polymerization with cytochalasin D (CytD)^38^, branched actin polymerization by Arp2/3 with CK666^39^, or linear actin polymerization by formin^40^ with SMIFH2^41^ in C2C12 cells at day two in DM (Fig. 3a,b). While all three drugs did not affect myoblast differentiation, myoblast fusion was significantly inhibited by CytD and CK666, but not by SMIFH2 (Fig. 3b-d), suggesting a role for branched, instead of linear, actin polymerization in mouse myoblast fusion. Since the Arp2/3 complex is activated by the actin NPFs, including WASP and WAVE, we first asked which NPFs were expressed in myoblasts. The mouse genome encodes multiple WASP and WAVE family members^42^. Among all family members examined, only N-WASP (and its interacting protein WIP2), WAVE2, as well as Arp2 and Arp3, were expressed in both C2C12 cells and satellite cells and at similar levels before and during differentiation (Fig. 3e and Extended Data Fig. 2a). We therefore focused our subsequent analyses on these actin regulators.

**Figure 3.**
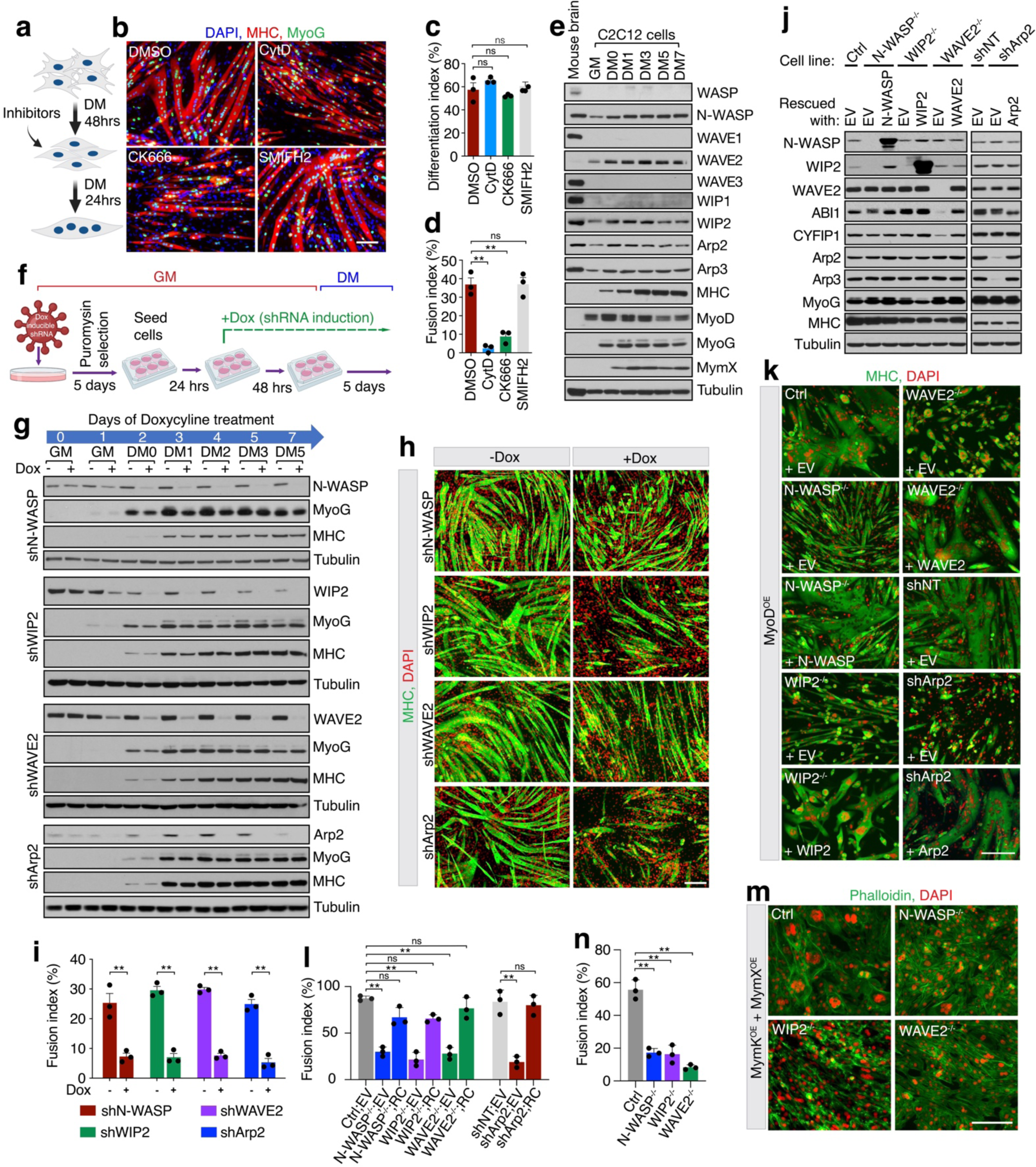
Role for branched actin polymerization in mouse myoblasts fusion. **(a)**Schematic diagram of pharmacological treatment of C2C12 cells with actin polymerization inhibitors. Upon 100% confluency, C2C12 cells were differentiated in DM for 48 hours, followed by incubation with DMSO (0.05%), a general actin polymerization inhibitor cytochalasin D (CytD; 18nM), Arp2/3 inhibitor (CK666; 50 μM) or formin inhibitor (SMIFH2; 10 μM) dissolved in DMSO for 24 hours. **(b)** Branched, but not linear, actin polymerization promoted C2C12 cell fusion. The DMSO or inhibitor treated cells shown in (**a**) were immunostained with anti-MyoG and anti-MHC at day three in DM (24 hours post actin polymerization inhibitor treatment). Note the severe fusion defects caused by CytD and CK666 treatment, but not by DMSO and SMIFH2. **(c-d)** The differentiation and fusion indexes of the cells shown in (**b**). **(e)** N-WASP, WAVE2 and WIP2 are highly expressed in C2C12 cells. C2C12 cells were grown to 100% confluency in GM then switched to DM. The cells were collected at the indicated time points during differentiation and subjected to western blotting for branched actin polymerization regulators, muscle differentiation markers and the muscle fusogenic protein myomixer (MymX). Adult mouse brain tissue lysate was used as the positive control for actin polymerization regulators. Note the high expression levels of N- WASP, WAVE2, and WIP2 among the homologous proteins. Three independent experiments were performed with similar results. **(f)** Schematic diagram of the conditional knockdown (KD) of the actin polymerization regulators during myoblast fusion. C2C12 cells were infected by lentivirus carrying doxycycline (Dox)-inducible shRNAs (Lenti-puromysin-Tet-On-shRNA-mCherry) against the target genes, followed by puromysin selection for five days. The surviving cells were plated at 60% in GM. After 24 hours, Dox or empty vehicle were added to GM. After 48 hours, the medium was switched to DM with Dox or empty vehicle for five days. **(g)** Western blots showing the conditional knockdown (KD) of the actin regulators during myoblast fusion. Control (-Dox) and KD (+Dox) cells as described in (**f**) were harvested at the indicated time points to examine the protein levels of NPFs, Arp2/3, and muscle differentiation markers (MyoG and MHC). Note the gradually decreased expression of target genes after the supplement of Dox, and that the expression of MyoG and MHC were comparable in control and KD cells throughout the differentiation process. **(h)** Conditional KD of the branched actin polymerization regulators caused server fusion defect. Immunostaining with anti-MHC and DAPI of the cells described in (**g**) at day five of differentiation. Note the sever fusion defects of the KD (+Dox) cells, compared to the control (-Dox) cells. **(i)** Quantification of the fusion index of control and KD cells shown in (**h**). **(j)** Depletion of N-WASP, WAVE2 or Arp2, but not WIP2, led to a decrease in protein expression of other complex members. Control (expressing a non-targeting sgRNA), KO, and KD cells were infected with either an empty retroviral vector (EV) or retroviruses containing rescue constructs (corresponding target genes with sgRNA- or shRNA-insensitive DNA cassettes). Three days after infection, cells were seeded at 60% confluence in GM. After 24 hours, the cells were infected by retroviruses containing MyoD. 48 hours post MyoD overexpression, the cells were collected and subjected to western blotting. Note that in N-WASP^-/-^ cells (EV lane), WIP2 level was significantly reduced; in WAVE2^-/-^ cells (EV lane), CYFIP1 and ABI1 levels were down; and in Arp2 KD cells (EV lane), Arp3 level was down. However, in WIP2^-/-^ cells (EV lane), N-WASP expression remained unchanged. In the rescued cells, expression of the complex members recovered to normal level. Three independent experiments were performed with similar results. **(k)** Immunostaining of cells describe in (**j**) with anti-MHC. Note the severe myoblast fusion defects in the KO and KD cells, and that overexpression of the corresponding genes rescued the myoblast fusion in the mutant cells. **(l)** Fusion indexes of cells shown in (**k**). **(m)** Knocking out NPFs inhibits myoblast fusion induced by myomaker (MymK) and myomixer (MymX) in GM. Control or NPF KO cells were seeded at 60% confluence in GM. After 24 hours, the cells were infected by either one retrovirus containing an empty vector (EV) or two retroviruses containing MymK and MymX (volume ratio 1:1), respectively. 24 hours post infection, the cells were fixed and stained with phalloidin. Note that control cells formed large multinucleated syncytia, but fusion of the KO cells was significantly inhibited. **(n)** Quantification of the fusion index of the cells shown in (**m**). Scale bars, 50 μm (**b**) and 200 μm (**h**, **k** and **m**). For (**c, d, i, l** and **n**), Mean ± s.d. values are shown in the bar graph, and significance was determined by two-tailed student’s t-test. Three independent experiments were performed. **: *p* < 0.01, ns: not significant.

Muscle differentiation involves two distinct but related processes – the expression of MRFs and muscle structural proteins, and the fusion between mononucleated myoblasts to generate multinucleated myofibers^12^. Inhibition of MRF expression could result in a defect and/or delay in myoblast fusion^43^. Since N-WASP, WIP2 and WAVE2 are expressed in muscle cells before and during differentiation, it is possible that they may affect both muscle-specific gene expression and myoblast fusion. Indeed, when we knocked down these genes in C2C12 cells several days before DM treatment, we observed a delay in MyoG expression, which could contribute in part to the severe myoblast fusion defect observed in these knockdown (KD) cells (Extended Data Fig. 2b- e). To investigate the specific functions of these actin regulators in myoblast fusion, we used the doxycycline (Dox)-inducible shRNA system to conditionally knock down these genes in C2C12 cells without affecting MyoG expression (Fig. 3f,g). Conditional KD of N-WASP, WIP2, WAVE2, or Arp2 all caused severe fusion defects, suggesting that these actin regulators are functionally required for myoblast fusion (Fig. 3h,i).

In addition to the KD experiments, we generated N-WASP, WIP2 and WAVE2 knockout (KO) C2C12 cell lines. Deleting WAVE2 did not affect the expression of N-WASP/WIP2, and vice versa (Fig. 3j). Interestingly, although deleting WIP2 did not affect the expression of N-WASP, N-WASP KO abolished WIP2 expression, suggesting that N-WASP is required to stabilize WIP2 (Fig. 3j). Since all the KO cells did not survive well in DM, we took advantage of the MyoD^OE^ system to analyze their fusion phenotypes in GM (Fig. 2a). As the conditional KD cells, the MyoD^OE^ KO cells also exhibited severe myoblast fusion defects despite normal MyoG and MHC expression. The fusion defects were rescued by expressing the corresponding knocked-out genes, demonstrating the specificity of the KOs (Fig. 3j-l). Since Arp2/3 KO cells were lethal, we generated Arp2 KD cells overexpressing MyoD in GM (Fig. 3j), which also exhibited a severe myoblast fusion defect (Fig. 3k,l). Moreover, it has been shown that co-expressing the fusogenic proteins MymK and MymX in C2C12 cells in GM induced robust cell fusion^31^. Knocking out N-WASP, WIP2, or WAVE2 significantly inhibited MymK/MymX-induced C2C12 cell fusion (Fig. 3m,n). Taken together, our loss-of-function analyses demonstrate critical functions for specific actin NPFs and interacting proteins in mammalian myoblast fusion.

### Distinct localization patterns of N-WASP/WIP2 and WAVE2 at the fusogenic synapse

Given the requirement for N-WASP, WIP2, WAVE2, and Arp2/3 in myoblast fusion, we asked whether these proteins are present at the fusogenic synapse. Live imaging of MyoD^OE^ C2C12 cells revealed strong enrichment of mScarleti (mScar)-tagged N-WASP, WIP2 and Arp2, but not WAVE2, with the F-actin-enriched structure at the fusogenic synapse prior to fusion, and their dissolution with F-actin when fusion was completed (Fig. 4a and Supplemental Video 6). Since it was surprising that WAVE2 was required for myoblast fusion, but mScar-WAVE2 was not detected at the fusogenic synapse, we stained MyoD^OE^ C2C12 cells co-expressing mScar-N-WASP and mNG-WIP2 with an anti-WAVE2 antibody (Extended Data Fig. 3a). Strikingly, WAVE2 was seen enriched in a very thin layer immediately underneath the plasma membrane at the leading edge of invasive structures (Fig. 4b), which was difficult to detect by live imaging. In contrast, N-WASP and WIP2 colocalized with the dense F-actin structure (Fig. 4b). Thus, N-WASP/WIP2 and WAVE2 exhibit distinct and complementary localization patterns during invasive protrusion formation in myoblast fusion.

**Figure 4.**
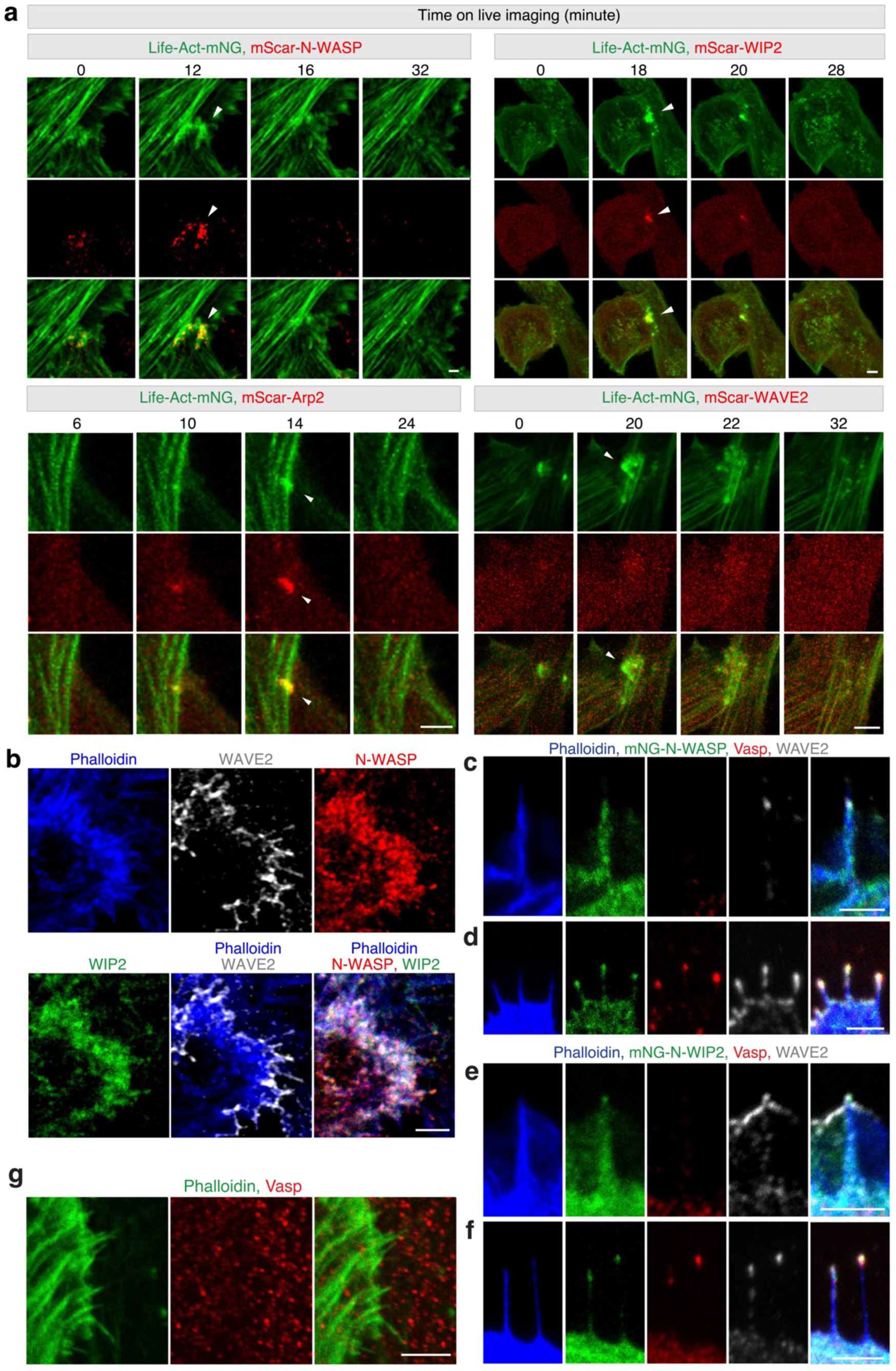
The localization of branched actin polymerization machinery in invasive protrusions. **(a)**Enrichment of N-WASP, WIP2 and Arp2 within the F-actin structure at the fusogenic synapse of C2C12 cells. Stable C2C12 cell lines co-expressing LifeAct-mNeongreen (mNG) and mScar-N-WASP (or mScar-WIP2, Arp2-mScar, mScar-WAVE2) were infected with retroviruses containing MyoD and imaged as described in Fig. 2c. Note the co-enrichment of F-actin and N-WASP, WIP2 and Arp2, but not WAVE2, at the fusogenic synapses (indicated by arrowheads) (see Supplemental Video 6). At least 15 fusogenic synapse were imaged for each target protein with similar results. **(b)** Complementary localization of WAVE2 and NWASP/WIP2 in the invasive structure. C2C12 cells co-expressing mNG-WIP2 and mScar-N-WASP were infected by retroviruses containing MyoD. After 48 hours, the cells were immunostained with anti-WAVE2. Note that WAVE2 was enriched at the leading edge of the invasive front as a thin layer underneath the plasma membrane, whereas N-WASP/WIP2 colocalized with F-actin within the invasive structure. *n* = 60 cells were imaged with similar results. **(c-f)** Invasive protrusions and filopodia are molecularly distinct. After 48 hours of MyoD overexpression, mNG-N-WASP or mNG-WIP2 expressing C2C12 cells were immunostained with anti-WAVE2, anti-Vasp and phalloidin. Note that the thick Vasp^−^ protrusions (**c** and **e**) had enriched N-WASP/WIP2 in the shafts, whereas the Vasp^+^ filopodia (**d** and **f**) contained Vasp and N-WASP/WIP2 enrichment at the tips. *n* = 75 protrusions of each type in at least 30 cells were imaged with similar results. **(g)** Vasp is not enriched at the tips of the invasive protrusions at the myoblast contact sites. After 48 hours of MyoD overexpression, wild-type C2C12 cells were immunostained with anti-Vasp and phalloidin. Note that Vasp was not enriched at the tips of the invasive protrusions. Scale bars, 5 μm (**a**), 3 μm (**b**, **e**, **f** and **g**) and 2 μm (**c** and **d**).

### Invasive protrusions are Ena/Vasp negative and show N-WASP/WIP2 enrichment in the shafts

The actin elongation factor Ena/Vasp has been shown to promote the growth of cellular protrusions, such as filopodia^44-46^. To ask whether invasive protrusions also employ Ena/Vasp for their elongation, we stained the MyoD^OE^ C2C12 cells with an antibody against Vasp, which is known to localize at the tips of filopodia^44-47^. Interestingly, both Vasp^−^ and Vasp^+^ protrusions were observed emanating from these cells (Extended Data Fig. 4a-c). The Vasp^−^ protrusions contained high level of N-WASP/WIP2 in the shafts (Fig. 4c,e), whereas in the Vasp^+^ protrusions, N-WASP/WIP2 were only enriched at the tips as Vasp (Fig. 4d,f). In both types of protrusions, WAVE2 was also localized at the tips, consistent with its close association with the plasma membrane (Fig. 4c-f). Morphologically, the Vasp^−^ protrusions appeared thicker and straighter, whereas the Vasp^+^ protrusions were thinner and often floppy (Extended Data Fig. 4b,c). The thicker and straighter Vasp^−^ protrusions enriched with N-WASP/WIP2 in the shafts are likely to be invasive, based on the previous work in *Drosophila* showing that the WASP complex is required for the invasiveness of protrusions^15^. Indeed, all the invasive protrusions at the muscle cell contact sites were Vasp^−^ (Fig. 4g). Taken together, we have distinguished invasive protrusions from filopodia based on their different molecular composition. We will refer to the Vasp^−^ protrusions with N-WASP/WIP2 enrichment in the shafts as invasive protrusions and the Vasp^+^ protrusions with no N-WASP/WIP2 enrichment in the shaft as filopodia hereafter.

### WAVE2 and N-WASP/WIP2 exhibit different functions in invasive protrusion formation

To investigate the roles for WAVE2 and N-WASP/WIP2 in generating invasive protrusions, we performed live imaging of wild-type and NPF KO C2C12 cells overexpressing MyoD. Wild-type cells exhibited dynamic actin cytoskeletal activities at the cell cortex, formation both lamellipodia and thick protrusions emanating from the lamellipodia (1± 0.26 new protrusion per minute per 10 μm of cell cortex, n = 90) (Fig. 5a,b and Supplemental Video 9). The thick protrusions were enriched with N-WASP/WIP2 in the shafts and should be invasive (Fig. 5c,d; Extended Data Fig. 3b,c; Supplemental Video 7,8). WAVE2^-/-^ cells barely generated any lamellipodia or finger-like protrusions (0.3 ± 0.13 new protrusion per minute per 10 μm of cell cortex, n = 60) (Fig. 5a,b and Supplemental Video 9), consistent with the previous finding that WAVE-mediated dendritic network in lamellipodia provides the source of actin filaments for nucleating finger-like protrusions^48^. Of the few protrusions observed in WAVE2^-/-^ cells, they were Vasp^−^ with N-WASP enrichment in the shafts, indicating that they would be invasive (Fig. 5e). In contrast, N-WASP^-/-^ cells, in which both N-WASP and WIP2 were absent (Fig. 3j), formed WAVE2-mediated lamellipodia and Vasp^+^/WAVE2^+^ thin protrusions (Fig. 5a,b,f and Supplemental Video 9), implying a role for N-WASP (and/or WIP2) in generating thick protrusions. Such a role for N-WASP was confirmed by treating MyoD^OE^ C2C12 cells with an N-WASP-specific inhibitor Wiskostatin^49^, which inhibited the formation of thick protrusions without affecting lamellipodia (Fig. 5h and Supplemental Video 10). However, WIP2^-/-^ cells (N-WASP was normal) (Fig. 3j), did generate many protrusions with N-WASP, WAVE2, and Vasp enriched at the tips, but they were bendy (Fig. 5a,b,g and Supplemental Video 9). Thus, WIP2 is required for recruiting N-WASP to the shaft of invasive protrusions and enhancing the mechanical stiffness of these protrusions. Finally, Arp2 KD cells exhibited no lamellipodia or finger-like protrusions, despite having normal cytosolic actin stress fibers (Fig. 5a,b and Supplemental Video 9). Taken together, these results suggest that branched actin polymerization is required for generating invasive protrusions and that WAVE2, N-WASP, and WIP2 have distinct functions in protrusion formation.

**Figure 5.**
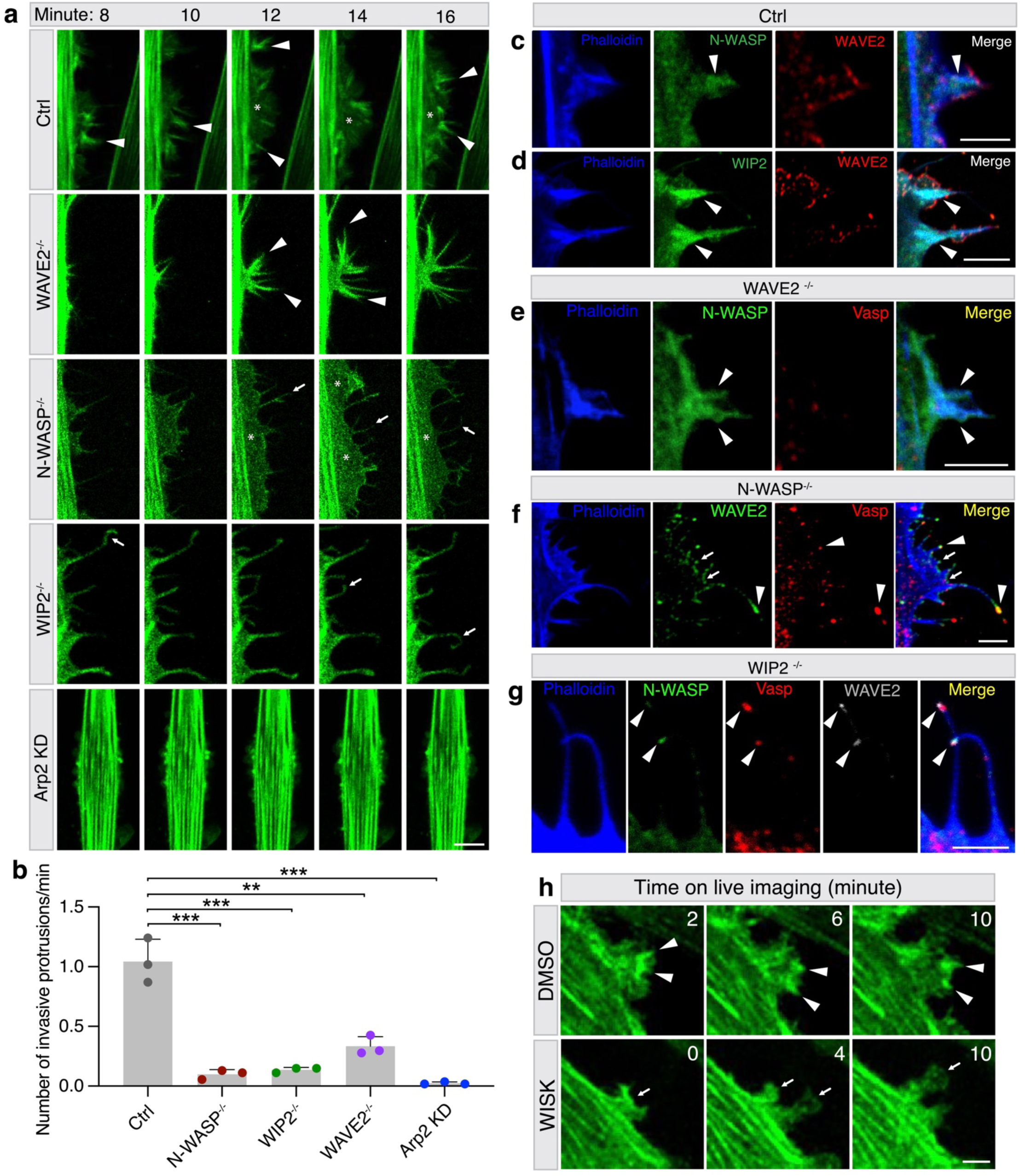
N-WASP/WIP2 and WAVE2 have different functions during protrusion formation. **(a)** Still images of cortical protrusions of wild-type, NPF KO and Arp2 KD C2C12 cells at 48 hours post MyoD overexpression. The N-WASP^-/-^, WIP2^-/-^, WAVE2^-/-^ and Arp2 KD cells expressing LifeAct-mNG were subjected to live imaging as described in **Fig.2c** (see Supplemental Video 9). Randomly picked straight/stiff protrusions (arrowheads), bendy/soft protrusions (arrows), and lamellipodia (asterisks) are shown. Note the straight/stiff vs. bendy/soft finger-like protrusions projected out of the protruding front of lamellipodia in wild-type vs. N-WASP^-/-^ cells; the absence of lamellipodia in WIP2^-/-^, WAVE2^-/-^, and Apr2-KD cells; the bendy/soft protrusions in WIP2^-/-^ cell; decreased number of protrusions in WAVE2^-/-^ cell; and no protrusions in Arp2 KD cells. **(b)** Quantification of the number of newly formed straight/stiff protrusion per 10 μm cell boundary per minute for cells with the genotypes shown in (**a**). *n* = 20-30 cells were measured for each genotype in each experiment and three independent experiments were performed. Mean ± s.d. values are shown in the bar graph, and significance was determined by two-tailed student’s t-test. **: *p* < 0.01. **(c-g)** Localization of N-WASP, WIP2, WAVE2 and Vasp in individual protrusions of wild-type and mutant MyoD^OE^ C2C12 cells. The wild-type (**c** and **d**), mNG-N-WASP expressing WAVE2^-/-^ (**e**), N-WASP^-/-^ (**f**), and mNG-N-WASP expressing WIP2^-/-^ (**g**) cells were immunostained with anti-WAVE2, phalloidin and/or Vasp at 48 hours post MyoD overexpression. Note that N-WASP and WIP2 (arrowheads) were enriched in the shafts of the invasive protrusions in wild-type and WAVE2^-/-^ cells, WAVE2 was enriched at the leading edge of the lamellipodia (arrows) including tips of protrusions where VASP was enriched (arrowheads) in N-WASP^-/-^ cells, and that N-WASP, WAVE2 and Vasp (arrowheads) were all enriched at the tips of the protrusions in WIP2^-/-^ cells. Three independent experiments were performed with similar results. **(h)** N-WASP is required for generating dynamic finger-like protrusions. At 48 hours post MyoD overexpression, 0.05% DMSO or 5 μM WISK dissolved in DMSO was added to the medium of the LifeAct-mNG expressing C2C12 cells and cells were immediately subjected to live imaging. The invasive protrusions are indicated by arrowheads, and lamellipodia by arrows, respectively (see Supplemental Video 10). Three independent experiments were performed with similar results. Scale bars: 10 μm (**a** and **h**), 3 μm (**c**, **e**, **f** and **g**) and 5 μm (**d**).

### WIP2 bundles F-actin and increases the mechanical strength of invasive protrusions

It is intriguing that in the absence of WIP2, N-WASP is relocated to the tips of bendy protrusions (Fig. 5g), indicating a crucial role for WIP2 in localizing N-WASP in the shafts and enhancing the mechanical strength of protrusions. To investigate the mechanism by which WIP2 functions in protrusions, we first examined WIP2-actin interaction using purified WIP2 N-terminal fragments. Previous studies identifying two actin-binding WH2 domains (WH2-1 and WH2-2) at the N-terminus of WIP proteins^19,50^. Using purified WH2-1, WH2-2, and WIP2-N (an N-terminal fragment containing both WH2-1 and WH2-2), we confirmed by high-speed co-sedimentation assays that both WH2 domains bound F-actin (Extended Data Fig. 5a). But to our surprise, low-speed co-sedimentation assays showed that WIP2-N (Fig. 6a), but not WH2-1 or WH2-2 alone (Extended Data Fig. 5b), bundled F-actin in a dosage-dependent manner. Thus, both WH2 domains are required for WIP2’s actin-bundling activity, suggesting that WIP2 functions as a two-filament actin crosslinker. Using total internal reflection fluorescence (TIRF) microscopy, we observed that when WIP2-N was added to a dynamically polymerizing branched actin network, it rapidly “froze” the network by forming actin bundles at numerous locations (Fig. 6b and Supplemental Video 11). At the single-filament level, WIP2 could induce bundling of two connected or unconnected actin filaments (Fig. 6c and Supplemental Video 12). Structured illumination microscopy (SIM) revealed that WIP2 appeared as sparsely localized patches along the actin bundles (Fig. 6d). Consistent with this, negative-stain EM also revealed light colored patches (formed by WIP2) along the tight actin bundles (Fig. 6e). Unlike the dynamin-mediated actin bundle, each of which is organized by a dynamin helix and has an average diameter of ∼34 nm^21^, the WIP2-mediated actin bundles did not have a fixed diameter and exhibited various sizes (Fig. 6e). Taken together, our data have demonstrated a novel function for WIP2 in bundling actin filaments via its two WH2 domains and a critical function for WIP2 in enhancing the mechanical strength of invasive protrusions, likely by bundling actin filaments in the shafts of the protrusions.

**Figure 6.**
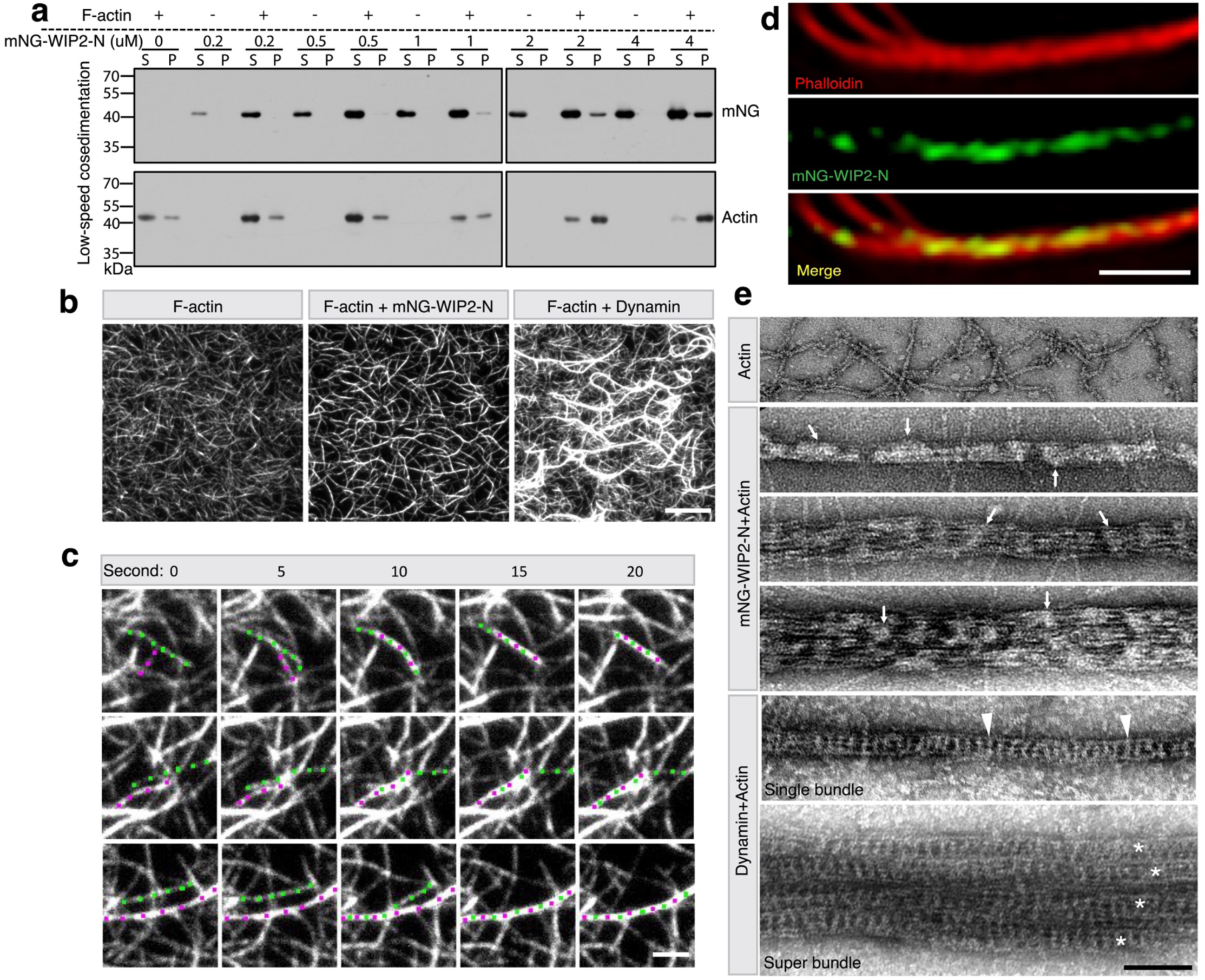
WIP2 bundles actin filaments. **(a)** WIP2 bundles actin filaments in a dose-dependent manner. 5 μM F-actin was incubated with increasing concentrations of mNG-WIP2-N. Supernatant (S) and pellet (P) were subjected to SDS-PAGE after low-speed centrifugation at 13,600g. Note the increasing amount of actin in the pellet with increasing concentrations of mNG-WIP2-N. **(b)** TIRF images of WIP2-mediated actin bundling. Arp2/3-mediated branched actin network was assembled prior to the addition of mNG-WIP2-N or Shibire (*Drosophila* dynamin) (see Supplemental Video 11). Note that the WIP2-mediated actin bundles were thinner than those of dynamin’s. Three independent imaging experiments were performed with similar results. **(c)** Still images of time-lapse TIRF imaging showing three examples of mNG-WIP2- mediated actin filament bundling, either between two connected actin filaments (top panel) or between two unconnected actin filaments (bottom two panels) (see Supplemental Video 12). The merging actin filaments are indicated by magenta or green dots. Three independent imaging experiments were performed with similar results. **(d)** Structure illumination microscopy (SIM) images of WIP2–actin bundles. Images were collected after a 30 minutes incubation of F-actin with alexa fluor 568-phalloidin (red) and mNG-WIP2-N (green). Three independent experiments were performed with similar results. **(e)** Electron micrographs of negatively stained actin filaments, WIP2-actin bundles, and dynamin-actin bundles. Note that in negatively stained samples, protein-enriched areas appear whitish. Three WIP2-actin bundles with different diameters are shown. A single dynamin-actin bundle and a dynamin-actin super bundle (each individual bundle is marked by an asterisk) are shown. Note the patchy distribution of mNG-WIP2-N (arrows indicate a few randomly selected mNG-WIP2-N patches) along the WIP2-actin bundles, compared to the helical structure of dynamin (arrowheads indicate a couple of randomly selected dynamin helical rungs). Three independent experiments were performed with similar results. Scale bars, 10 μm (**b**), 2 μm (**c**), 1 μm (**d**) and 100 nm (**e**).

### Dynamin is a “pioneer” actin-bundling protein during invasive protrusion growth

Now that we have identified a crucial role for Arp2/3 and its NPFs in invasive protrusion formation, we asked how newly polymerized branched actin filaments are organized into bundles. To observe the dynamics of nascent actin polymerization during protrusion formation, we first performed live imaging of MyoD^OE^ C2C12 cells expressing LifeAct-mNG and Arp2-mScar (Supplemental Video 13). Strikingly, we observed low-density actin “clouds” growing out of the side of pre-existing F-actin bundles within a protrusion (Fig. 7a, minute 3-5). The actin cloud had Arp2 enrichment at the expanding front, indicating that it contained newly polymerized branched actin filaments. Shortly after, new F-actin bundles arose from within the actin cloud, coinciding with a decrease of Arp2 level in the bundles. These new bundles, in turn, provided additional sites from which new Arp2/3-mediated actin clouds could emerge (Fig. 7a, minute 7).

**Figure 7.**
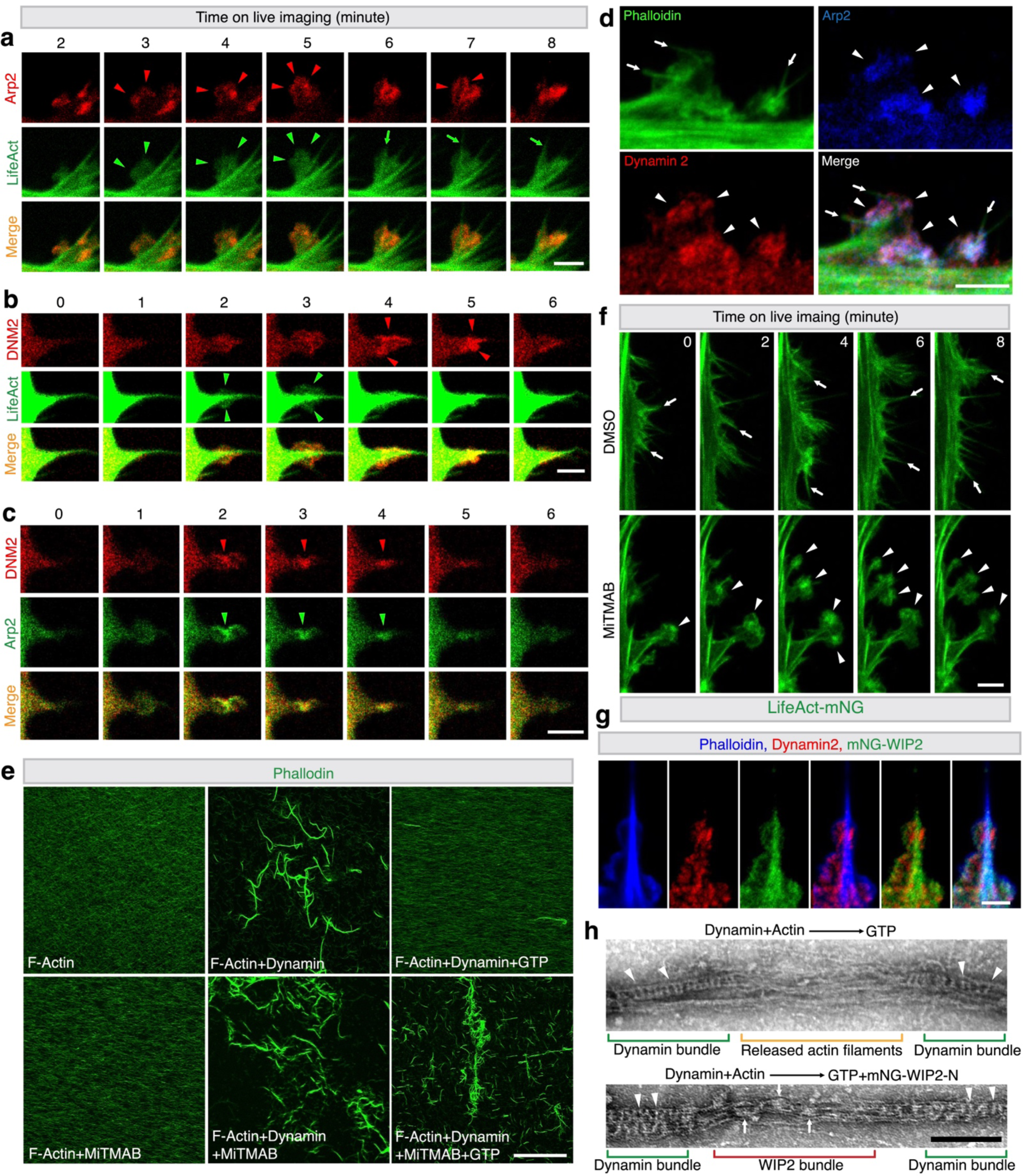
Dynamin and WIP2 cooperatively bundle newly formed actin filaments nucleated by the Arp2/3 complex to form mechanically stiff bundles. **(a-c)** Dynamic enrichment of Arp2 and dynamin with newly polymerized branch actin filaments. C2C12 cells co-expressing LifeAct-mNG/Arp2-mScar (**a**), LifeAct-mNG/mScar-dynamin 2 (**b**), or Arp2-mNG/mScar-dynamin 2 (**c**) were infected with retrovirus containing MyoD. 48 hours post infection, the cells were subjected to live imaging (see Supplemental Video 13-15). Note that newly polymerized branched actin appeared as actin “clouds” emanating from the sides of the actin bundles, and that Arp2 and dynamin 2 were both co-enriched with these new actin clouds. In (**a**), red arrowheads point to enriched Arp2-mScar at the leading front of branched actin polymerization, and the resulting branched actin-containing actin clouds are indicated by green arrowheads. An emerging F-actin bundle from the actin cloud is indicated by a green arrow. In (**b**), red arrowheads point to mScar-dynamin 2 colocalizing with the actin clouds containing nascent branched actin filaments (green arrowheads) (2 and 3 min). In 4 and 5 min, mScar-dynamin 2 was enriched along the protrusion, bringing the newly polymerized branched actin filaments into the main F-actin bundle. In (**c**), the co-enriched mScar-dynamin 2 and Arp2-mNG are indicated by red and green arrowheads, respectively. Three independent experiments were performed with similar results. **(d)** Triple labeling of fixed C2C12 cells by Arp2-mNG, anti-dynamin 2 and phalloidin. C2C12 cells expressing Arp2-mNG were infected with retrovirus containing MyoD. After 48 hours, cells were immunostained with anti-dynamin 2 and phalloidin. Note the co-localization of Arp2 and dynamin 2 with the actin clouds (indicated by arrowheads) at the periphery of actin bundles (indicated by arrows). Three independent experiments were performed with similar results. **(e)** Dynamin inhibitor MiTMAB inhibited dynamin helix disassembly in the presence of GTP. F-actin pre-assembled from 1 μM G-actin was incubated with or without 1 μM dynamin for 30 minutes at RT. Subsequently, 2mM GTP with or without 40 μM MiTMAB was added to the appropriate reaction mixtures and incubated for five minutes. The samples were then mixed with alexa fluor 488-conjugated phalloidin and subjected to confocal imaging on glass coverslips. Note that the dynamin-induced actin bundles were disassembled by GTP addition (top right panel). MiTMAB neither bundled actin by itself (bottom left panel) nor inhibited dynamin-induced actin bundling (bottom middle panel), but it inhibited actin bundle disassembly in the presence of GTP (bottom right panel). **(f)** Dynamic disassembly of dynamin helices is essential for proper actin bundle formation in the shafts of invasive protrusions. The LifeAct-mNG-expressing C2C12 cells were infected with retrovirus containing MyoD. 24 hours post infection, empty vehicle (0.05% DMSO) or 5 μM MiTMAB dissolved in DMSO was added to the GM. After another 24 hours, the cells were subjected to live imaging (see Supplemental Video 16). Still images of cortical protrusions are shown. Note that the control cells (top panels) generated tightly organized actin bundles (arrowheads), whereas the MiTMAB- treated cells (bottom) had persistent actin clouds (arrows) that were not brought into the main bundles. Three independent experiments were performed with similar results. **(g)** Distinct localization of WIP2 and dynamin 2 along the invasive protrusions. C2C12 cells expressing mNG-WIP2 were immunostained with anti-dynamin 2 and phalloidin at 48 hours post MyoD overexpression. Note that although both WIP2 and dynamin 2 were observed in F-actin clouds at the periphery of the shaft, WIP2, but not dynamin 2, was highly enriched within the shaft of the protrusion. Three independent experiments were performed with similar results. **(h)** WIP2 stabilizes actin bundles after dynamin helix disassembly. F-actin pre-assembled from 1 μM G-actin was incubated with 1 μM dynamin for 30 minutes to form multi-filament bundles. These bundles were then incubated with 1mM GTP without (top) or with mNG-WIP2-N (bottom panel) for 30 minutes, and subjected to negative staining and electron microscopy. Arrowheads and arrows indicate several randomly picked dynamin helical rungs and mNG-WIP2-N patches, respectively. Note that GTP addition could result in partial disassembly of dynamin helices and loosening of the dynamin-mediated actin bundles, and that mNG-WIP2-N bound the loosened actin filaments to tighten the bundle. Sale bars, 3 μm (**a-c**), 4 μm (**d**), 50 μm (**e**), 5 μm (**f**), 2 μm (**g**) and 200 nm (**h**).

To investigate which factor(s) are involved in organizing nascent branched actin filaments in the low-density actin clouds into bundles, we examined the localization of dynamin, which is a potent multi-filament actin-bundling protein required for invasive protrusions formation at the *Drosophila* fusogenic synapse^21^. Live imaging in MyoD^OE^ C2C12 cells co-expressing mScar-dynamin 2 and LifeAct-mNG revealed striking colocalization between dynamin 2 and the F-actin clouds at the initial stage of branched actin polymerization (Fig. 7b, minute 2 and 3 and Supplemental Video 14). Subsequently, dynamin 2 was transiently enriched on the periphery of the actin bundle, coinciding with the thickening of the protrusion (Fig. 7b, minute 4 and 5), indicating that dynamin 2 was involved in crosslinking the new actin filaments into the pre-existing actin bundles. Live imaging of MyoD^OE^ C2C12 cells co-expressing mScar-dynamin 2 and Arp2-mNG revealed that mScar-dynamin 2 colocalized with Arp2 (Fig. 7c, minute 2- 4 and Supplemental Video 15). Moreover, immunostaining with an anti-dynamin 2 antibody showed that the endogenous dynamin 2 also co-localized with Arp2 and the F- actin clouds in Arp2-mNG-expressing cells (Fig. 7d). These observations strongly suggest that dynamin functions as a “pioneer” actin-bundling protein that organizes the nascent branched actin filaments into pre-existing actin bundles during protrusion growth.

If dynamin functions as a pioneer actin-bundling protein, one would expect that disrupting dynamin function would disrupt actin bundle formation in the protrusions. To test this, we took advantage of a cell-permeable dynamin inhibitor, MiTMAB, which inhibits dynamin GTPase activity by targeting the PH domain^51^. Our previous study demonstrated that dynamin bundles actin during its assembly into helical structures, followed by rapid disassembly of the dynamin helices upon GTP hydrolysis, leaving behind loosely bundled actin filaments^21^. The PH domain is not required for actin bundling, but for disassembly of the dynamin helix in the presence of GTP^21^. We therefore tested the effect of MiTMAB in dynamin-mediated actin bundling *in vitro*. In the presence of MiTMAB, the dynamin-actin bundles formed normally but failed to disassemble in the presence of GTP (Fig. 7e). Thus, MiTMAB inhibited the dynamic cycles of dynamin-mediated actin bundling and GTP hydrolysis-triggered dynamin helix disassembly. Correspondingly, treating MyoD^OE^ C2C12 cells with MiTMAB resulted in persistent actin clouds along the protrusions, compared to the properly formed long and narrow protrusions in control cells (Fig. 7f and Supplemental Video 16). Thus, MiTMAB “froze” the dynamin-actin bundles in the actin clouds and disrupted the dynamic process of actin bundling required for invasive protrusion formation. Taken together, we conclude that dynamin is a key actin-bundling protein required for bringing nascent branched actin filaments into preexisting bundles during invasive protrusion growth.

### WIP2 stabilizes the dynamin-mediated actin bundles

It is intriguing that dynamin 2 largely colocalized with F-actin clouds instead of with the shafts of the protrusions, revealed by immunostaining with anti-dynamin 2 in mNG- WIP2-expressing cells (Fig. 7g). How to stabilize the loose actin bundles following dynamin helix disassembly in the shaft was unclear. Given WIP2’s ability to bundle actin (Fig. 6a,b), we asked whether WIP2 could be involved in bundle stabilization. Indeed, compared to its presence in the actin clouds, WIP2 was more strongly enriched with the actin bundles in the shafts (Fig. 7g). Since live imaging experiments demonstrated that the F-actin clouds preceded actin bundle formation (Fig. 7a), the distinct spatial localization patterns of dynamin 2 and WIP2 strongly suggest that WIP2 stabilizes the actin bundles in the shaft following dynamin disassembly. In support of this, negative staining experiments revealed that, upon GTP addition, stretches of loose actin filaments that were no longer associated with dynamin 2 could be further crosslinked into tight bundles by WIP2 (Fig. 7h).

### Arp2/3-mediated actin polymerization promotes invasive protrusion formation and fusion of mouse myoblasts *in vivo*

We next asked whether branched actin polymerization is required for invasive protrusion formation and myoblast fusion *in vivo*. Using the MyoD^iCre^ driver line^52^, we specifically knocked out ArpC2, a subunit of the Arp2/3 complex, in the embryonic muscle progenitors. The mutant embryos exhibited normal gross morphology and body size at the early developmental stage (E12.5) but had curved back and thinner limbs at later stages (E15.5 and 17.5), indicating that skeletal muscle development may be defective (Fig. 8a). In support of this, the bulk size of the limb muscles in E17.5 mutant embryos was significantly reduced compared to the control, shown by whole mount limb staining with anti-MHC (Fig. 8b). Consistently, the cross-section area and myofiber number of the limb muscles in E17.5 mutant embryos were also significantly reduced (Fig. 8c-e). On the other hand, the cell proliferation marker (Ki67) and myogenic/muscle structural proteins (MyoG and MHC) showed similar expression in the limb muscles between mutant and control embryos (Fig. 8f and Extended Data Fig. 6). Therefore, we concluded that the significant decrease in the muscle size in ArpC2 mutant embryos is not due to defects in myoblast proliferation or myogenic/structural gene expression, but due to defects in myoblast fusion. In support of this, immunostaining of the longitudinal limb sections of E17.5 mutant embryos with anti-MHC antibody revealed many differentiated (MHC^+^), but mononucleated myoblasts associated with elongated myofibers. In contrast, few mononucleated myoblasts were present in the control embryos at this stage (Fig. 8 g,h). Furthermore, TEM analysis showed that muscle cells in E15.5 mutant embryos seldom formed invasive protrusions at muscle cell contact sites (3.5% cells exhibited any protrusion; n = 57) compared to control embryos (30% cells exhibited protrusions; n = 30) (Fig. 8i). Taken together, these results demonstrate that Arp2/3-mediated branched actin polymerization is required for generating invasive protrusions to promote myoblast fusion *in vivo*.

**Figure 8.**
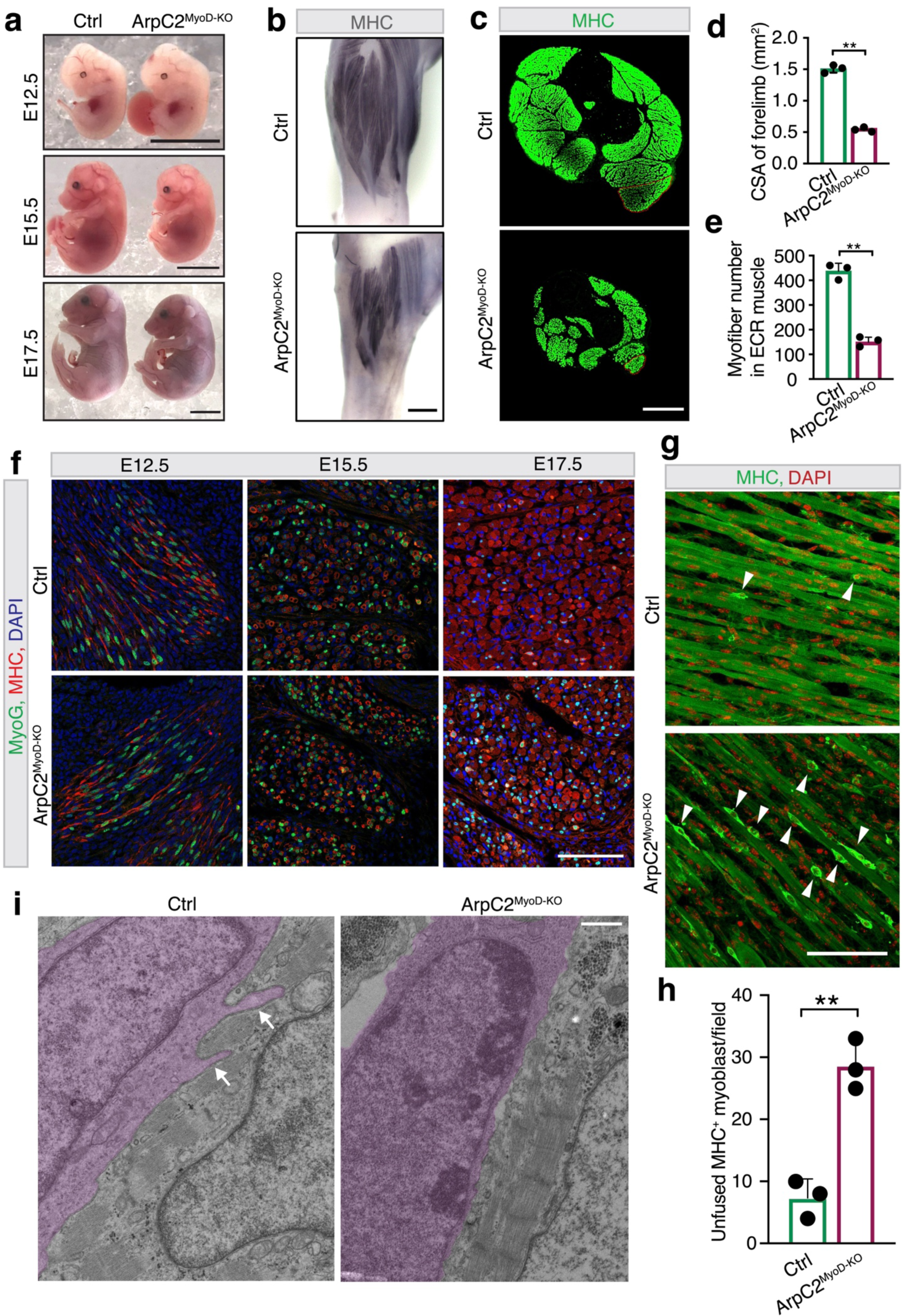
Arp2/3-mediated branched actin polymerization promotes invasiveprotrusion formation and fusion of mouse myoblasts *in vivo*. **(a)** Gross morphology of control and ArpC2^MyoD-KO^ embryos at E12.5, E15.5, and E17.5, respectively. ArpC2^MyoD-KO^ embryos exhibited normal size and morphology at early stage (E12.5), but abnormal spinal curvature and thinner limbs at late stages (E15.5 and E17.5). **(b)** Whole mount immunostaining with anti-MHC of forelimbs of control and ArpC2^MyoD-KO^ embryos at E17.5. **(c)** Forelimb cross section immunostaining with anti-MHC of control and ArpC2^MyoD-KO^ embryos at E17.5. The extensor carpi radialis longus (ECR) muscle is outlined. **(d)** Cross section area (CSA) of the forelimb skeletal muscle in control and ArpC2^MyoD-KO^ embryos shown in (c). Three control and ArpC2^MyoD-KO^ embryos were examined. **(e)** The number of myofibers in the ECR muscle of forelimbs in control and ArpC2^MyoD-KO^ embryos shown in (c). Three control and ArpC2^MyoD-KO^ embryos were examined. **(f)** Immunostaining for MyoG and MHC in the forelimb sections of control and ArpC2^MyoD-KO^ embryos at E12.5, E15.5 and E17.5. **(g)** Immunostaining for MHC in longitudinal sections of forelimbs in control and ArpC2^MyoD-KO^ embryos at E17.5. Randomly selected mononucleated MHC^+^ myoblasts are indicated by arrowheads. **(h)** Quantification of the number of mononucleated MHC^+^ myoblasts shown in (g). Three control and ArpC2^MyoD-KO^ embryos were examined, respectively. Mononucleated myoblasts in 12 40x microscopic fields of each embryo were counted. **(i)** TEM analysis of the forelimb muscle of control and ArpC2^MyoD-KO^ embryos at E15.5. The invading fusion partner is pseudo-colored in magenta. The invasive protrusions are indicated by arrows. Nine out of 30 (30%) muscle cells in the control embryos and two out of 57 (3.5%) muscle cells in the ArpC2^MyoD-KO^ embryos were observed having invasive protrusions. For (**d, e** and **h**), mean ± s.d. values are shown in the graph, and significance was determined by two-tailed student’s t-test. **: *p* < 0.01. Scale bars: 5 mm (**a**), 0.3 mm (**b**), 0.5 mm (**c**), 100 μm (**f** and **g**), and 1 μm (**i**).

## Discussion

In this study, we identified the molecular components of the asymmetric fusogenic synapse in mammalian myoblast fusion and uncovered the mechanisms underlying the formation of branched actin-driven invasive protrusions (Extended Data Fig. 7). We found that two Arp2/3 NPFs, WAVE2 and N-WASP, exhibit distinct localization patterns in invasive protrusions. While WAVE2 is closely associated with the plasma membrane at the leading edge of the protrusive structures, N-WASP is recruited by WIP2 to the actin bundles in the shafts of the protrusions. During protrusion growth, Arp2/3 nucleates branched actin filaments to generate low-density actin clouds at the periphery of a protrusion, accompanied by an initial round of bundling of the nascent actin filaments by dynamin, followed by WIP2-mediated actin bundle stabilization. Our findings revealed, for the first time, the temporal and spatial coordination of actin NPFs (WAVE2 and N-WASP) for branched actin polymerization and actin-bundling proteins (dynamin 2 and WIP2) in driving invasive protrusion formation.

How N-WASP and WAVE exert different functions in cellular protrusion formation is a longstanding question. In this study, we demonstrate that their different and complementary localization patterns in the protrusions are key to their distinct functions. Activation of Arp2/3 by WAVE at the leading edge of the invasive front provides a dendritic actin network where invasive protrusions will arise. On the other hand, activation of Arp2/3 by N-WASP along the actin bundles in the shaft provides additional branched actin filaments to thicken the bundles. Moreover, N-WASP’s shaft localization is dependent on WIP, the latter of which functions as a WH2 domain-mediated actin-bundling protein. This new insight on WIP highlights an essential role for WIP in imparting invasiveness of a protrusion and provides a logic explanation for the compromised protrusions in *sltr* (*WIP*) mutant in *Drosophila*^15^, the bendy protrusions in WIP2^-/-^ mouse myoblasts (Fig. 5a,g), and the lack of mechanically stiff actin core in podosomes and invadopodia in WIP^-/-^ osteoclasts and metastatic carcinoma cells^53,54^.

A prevailing view on how cellular protrusions form involves linear actin polymerization/elongation from a branched actin network within lamellipodia. It remained unclear whether branched actin filaments could be organized into long and narrow bundles to push the plasma membrane forward, and if this is possible, what are the underlying mechanisms. We have demonstrated here that the large GTPase dynamin is a key factor that organizes branched actin filaments into bundles. Our previous study has shown that dynamin is a highly efficient, dynamic actin-bundling protein^21^, but how dynamin spatially and temporally bundles actin filaments during invasive protrusion formation was unknown. In this study, we revealed that dynamin serves as a “pioneer” actin-bundling protein that organizes nascent branched actin filaments into bundles to facilitate protrusion formation and growth. Once dynamin forms a full helix that has bundled 12 or 16 actin filaments, it will be disassembled rapidly by GTP hydrolysis, freeing dynamin dimers for new rounds of actin bundling^21^. Consistent with this, dynamin is no longer enriched in well-formed actin bundles, the latter of which are stabilized by additional actin crosslinkers, such as WIP.

Our analyses of invasive protrusions in this study have clearly distinguished them from filopodia, even though both are actin-rich finger-like cellular protrusions. It is well known that invasive protrusions are mechanically stiff protrusions used by cells to drill their neighbors or basement membrane. The mechanical forces exerted by invasive protrusions are manifested by their ability to trigger mechanosensory responses in the receiving fusion partner^55,56^. In contrast, filopodia are thin and floppy protrusions that function as antennae for cells to probe their environment^47^. Although both types of protrusions arise from the cortical dendritic network formed by WAVE and Arp2/3^48,57^ (and this study), their growth/extension is driven by different actin nucleators and bundling proteins. The extension of filopodia depends on linear actin elongation factors, such as formin and Ena/Vasp, and the linear actin filaments are bundled by the actin crosslinker Fascin^47,58^. In contrast, we demonstrate in this study that the growth of invasive protrusions is driven by branched actin nucleator Arp2/3 and its NPFs, N- WASP and WAVE, and the branched actin filaments are bundled by dynamin and WIP. Because short, branched actin filaments are more suitable for doing mechanical work than long and bendy filaments^59^, invasive protrusions are Vasp^−^ and driven by N- WASP/WIP, whereas the Vasp^+^ filopodia are not invasive. It is worth noting that mechanically stiff, N-WASP/WIP^+^ protrusions are not unique to cell-cell fusion. Given that branched actin polymerization is required for the formation of other mechanically stiff protrusions, such as podosomes and invadopodia in cancer metastasis, leukocyte diapedesis, immunological synapse maturation, and uterine-vulval attachment^8,53,54,60-62^, it is conceivable that similar molecular mechanisms would underlie the formation of invasive protrusions in other cellular contexts beyond cell-cell fusion.

---

**Extended Data Figure 1.**
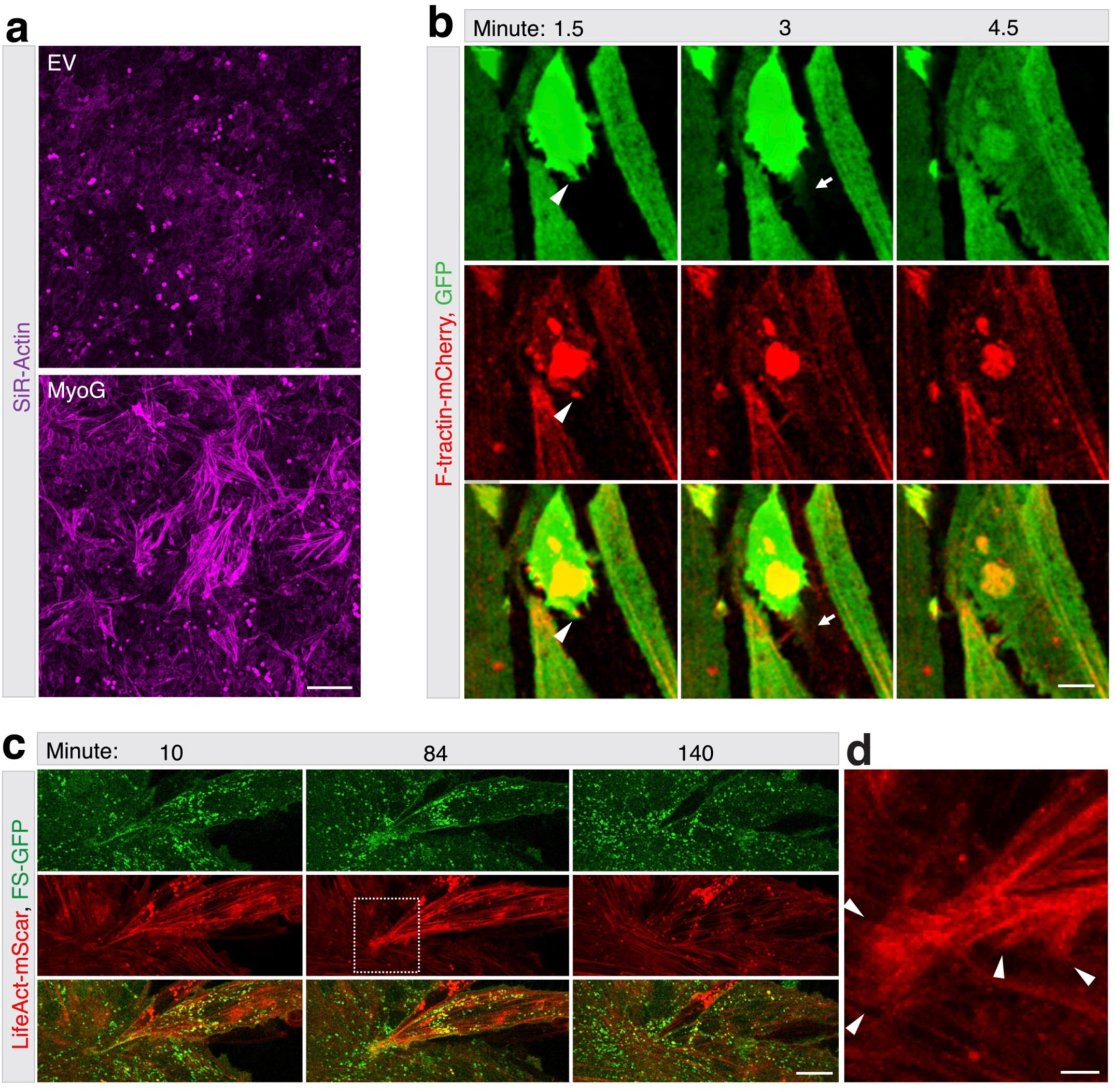
Myogenic transcription factors activate actin polymerization in heterologous cells and actin polymerization is dynamic at the fusion sites. **(a)**MyoG expression increased F-actin level in U2OS cells. U2OS cells were infected by lentivirus containing MyoG. Three days post-infection, the cells were incubated with SiR-Actin in culture medium for 30 minutes and fixed with 4% PFA. Note the increase in the F-actin level in MyoG-expressing cells (bottom panel) compared with the control with empty vector (EV) (top panel). Three independent experiments were performed with similar results. **(b)** Still images of a fusion event between C2C12 cells expressing F-tractin-mCherry. The F-tractin-mCherry expressing C2C12 cells with or without cytosolic GFP were co-cultured, switched to DM, and subjected to live imaging as described in **Fig.1d**. The arrowhead indicates the F-actin enriched membrane protrusion (1.5 min) that led to fusion pore formation and transfer of GFP from the top to the bottom cell (3 minutes, indicated by an arrow) (see Supplemental Video 3). 23 fusion events were observed with similar results. **(c-d)** Still images of a fusion event between mouse satellite cells. The satellite cells co-expressing F-tractin-mCherry and FS-GFP were cultures in DM for 24 hours, followed by live imaging. Boxed area is enlarged in (**d**). Note the actin enrichment indicated by arrowheads at 86 minutes prior to fusion (see Supplemental Video 4). 21 fusion events were observed with similar results. Scale bars: 200 μm (**a**), 10 μm (**b** and **c**) and 2 μm (**d**).

**Extended Data Figure 2.**
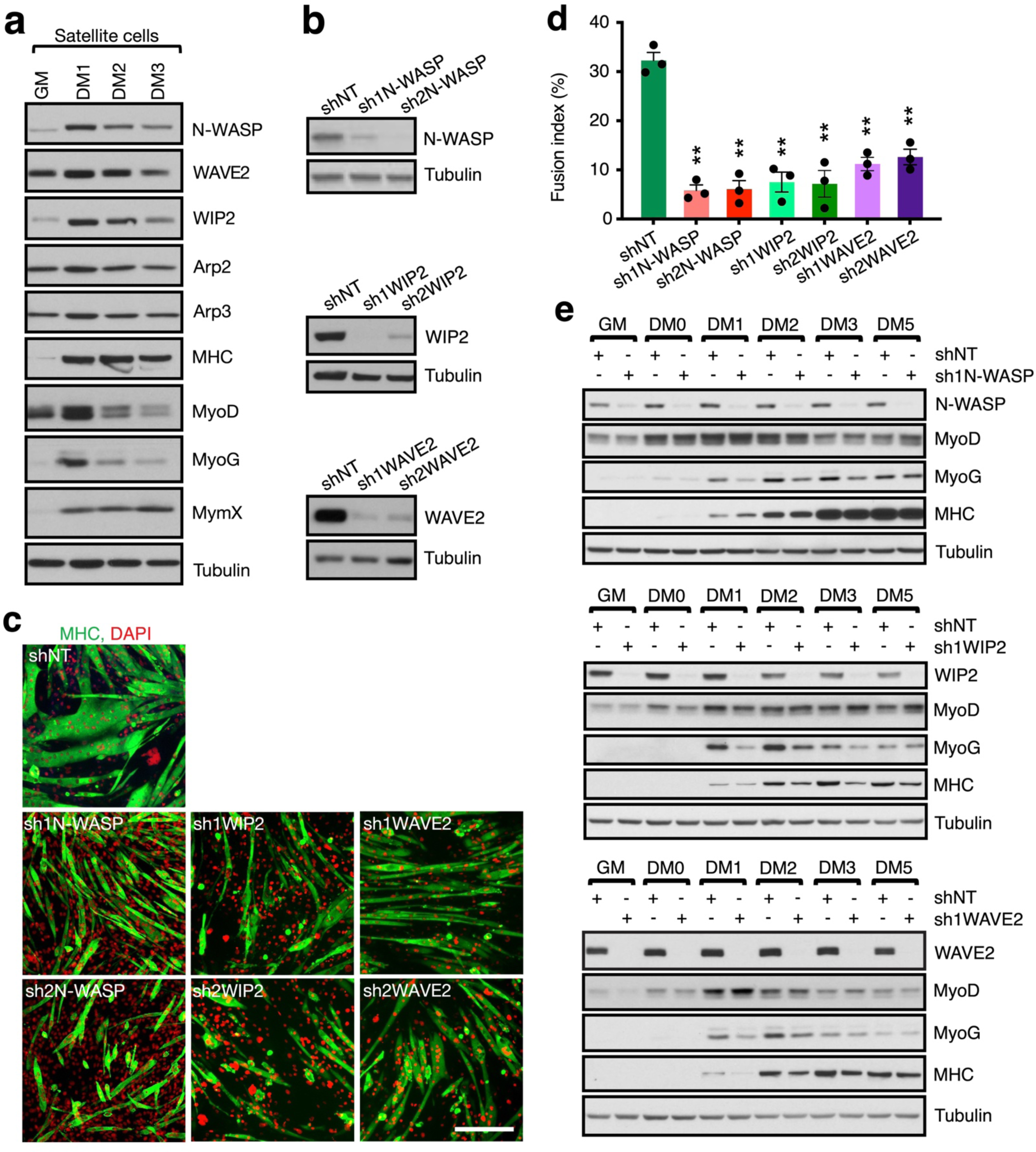
MyoG expression is delayed during differentiation when knocking down N-WASP, WIP2 and WAVE2 in C2C12 cells early during proliferation. **(a)**N-WASP, WAVE2 and WIP2 are highly expressed in mouse satellite cells. Mouse satellite cells were grown to 70% confluence in GM and switched to DM. The cells were collected at the indicated time points during differentiation and subjected to western blotting for NPFs, Arp2/3, muscle differentiation markers and the muscle fusogenic protein MymX. Three independent experiments were performed with similar results. **(b)** The knockdown (KD) efficiency of different shRNAs against target genes. C2C12 cells were infected by lentivirus (Lenti-shRNA-mCherry-puromysin) carrying a scramble shRNA (shNT) or shRNAs against the genes indicated. Two days later, cells underwent puromycin selection for five days. Then, the viable cells in GM were harvested for western blot. Two shRNAs were used to knock down each gene. Note the significant KD level of each target gene by both shRNAs. Three independent experiments were performed with similar results. **(c)** C2C12 cell fusion was significantly inhibited by knocking down N-WASP, WIP2 and WAVE2 for seven days before differentiation. C2C12 cells were treated with the shRNAs described in (**b**) in GM, and underwent puromycin selection for five days. Once the cell reached 100% confluence after one day, the GM was replaced with DM. At day five of differentiation, the cells were immunostained with anti-MHC. Scale bar, 50 μm. **(d)** Quantification of the fusion index of C2C12 cells shown in (**c**). Three independent experiments were performed. Mean ± s.d. values are shown in the bar graph, and significance was determined by two-tailed student’s t-test. **: *p* < 0.01 (compared to control). **(e)** Western blots of the NPFs and muscle differentiation markers of the control and NPF KD cells described in (**c**). The control (shNT) and NPF KD C212 cells were collected at the indicated time points and subjected to western blotting. Note that in DM1, DM2, and DM3, MyoG was expressed at a lower level in KD cells compared to the control. Three independent experiments were performed with similar results.

**Extended Data Figure 3.**
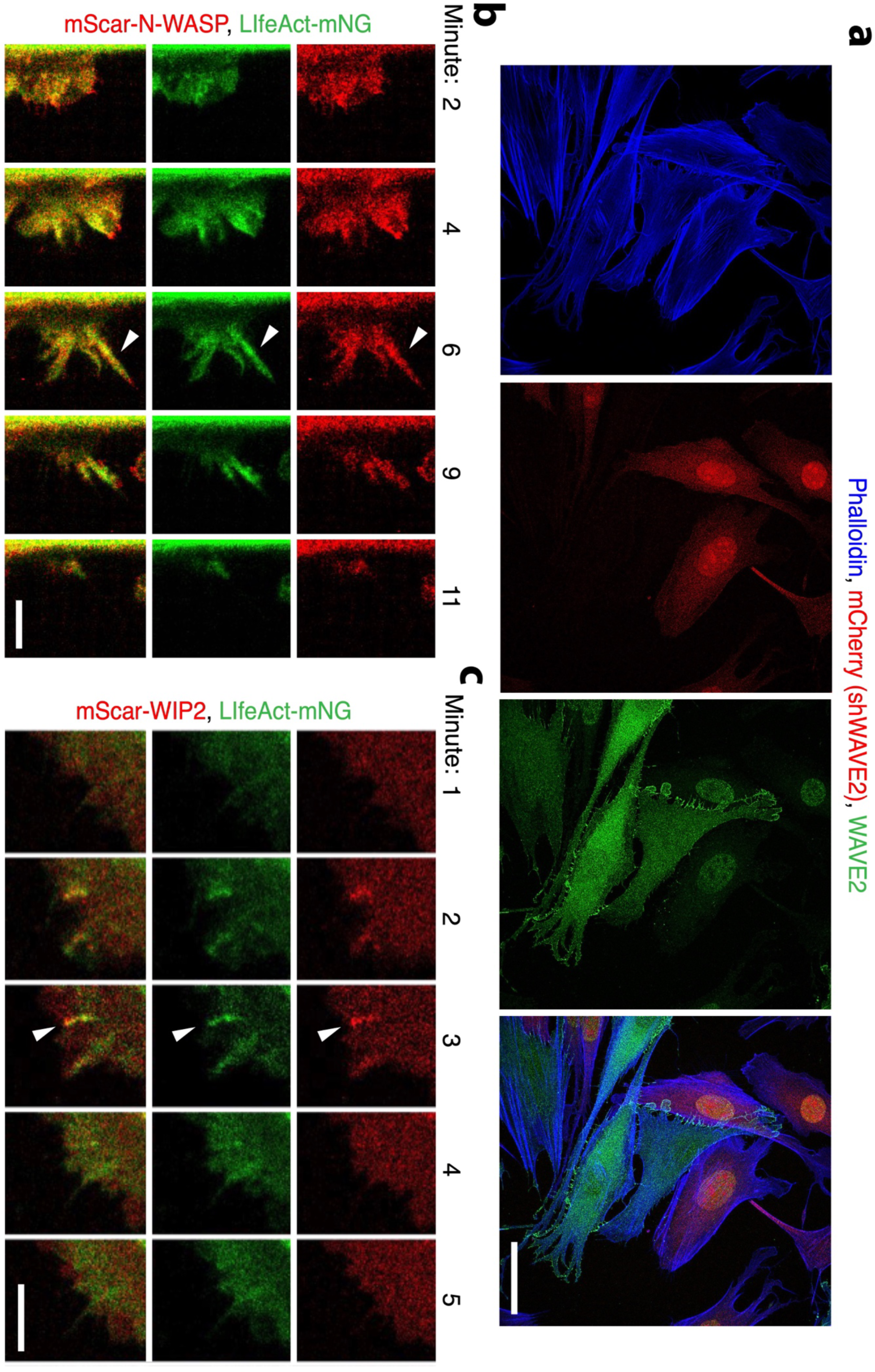
Dynamic enrichment of N-WASP and WIP2 in the shafts of straight/stiff protrusions. **(a)**Testing the specificity of the WAVE2 polyclonal antibody used in this study. The WAVE2 KD C2C12 cells were generated as described in **Extended Data Fig. 2b**. After five days of puromycin selection, the surviving cells were mixed with wild-type C2C12 cells and seeded at 30% confluence in GM. After 24 hours, the cells were immunostained with anti-WAVE2 and phalloidin. The WAVE2 KD cells were marked by mCherry expression. Note that the wild-type cells had enriched WAVE2 along the plasma membrane at cell-cell contact sites. The nuclear staining in WAVE2 KD cells is non-specific indicated by western blot (**Extended Data Fig. 2e**). Three experiments were performed with similar results. **(b-c)** Stills from a time-lapse series showing that N-WASP (**b**) and WIP2 (**c**) colocalize with F-actin along the entire length of straight protrusions. C2C12 cells co-expressing LifeAct-mNG and mScar-N-WASP (or mScar-WIP2) were infected with retrovirus containing MyoD. At 48 hours post MyoD overexpression, the cells were subjected to live imaging (see Supplemental Video 7 and 8). Arrowheads point to randomly selected protrusions. Three experiments for mScar-N-WASP and mScar-WIP2 each were performed with similar results. Scale bars: 30 μm (**a**) and 5 μm (**b** and **c**).

**Extended Data Figure 4.**
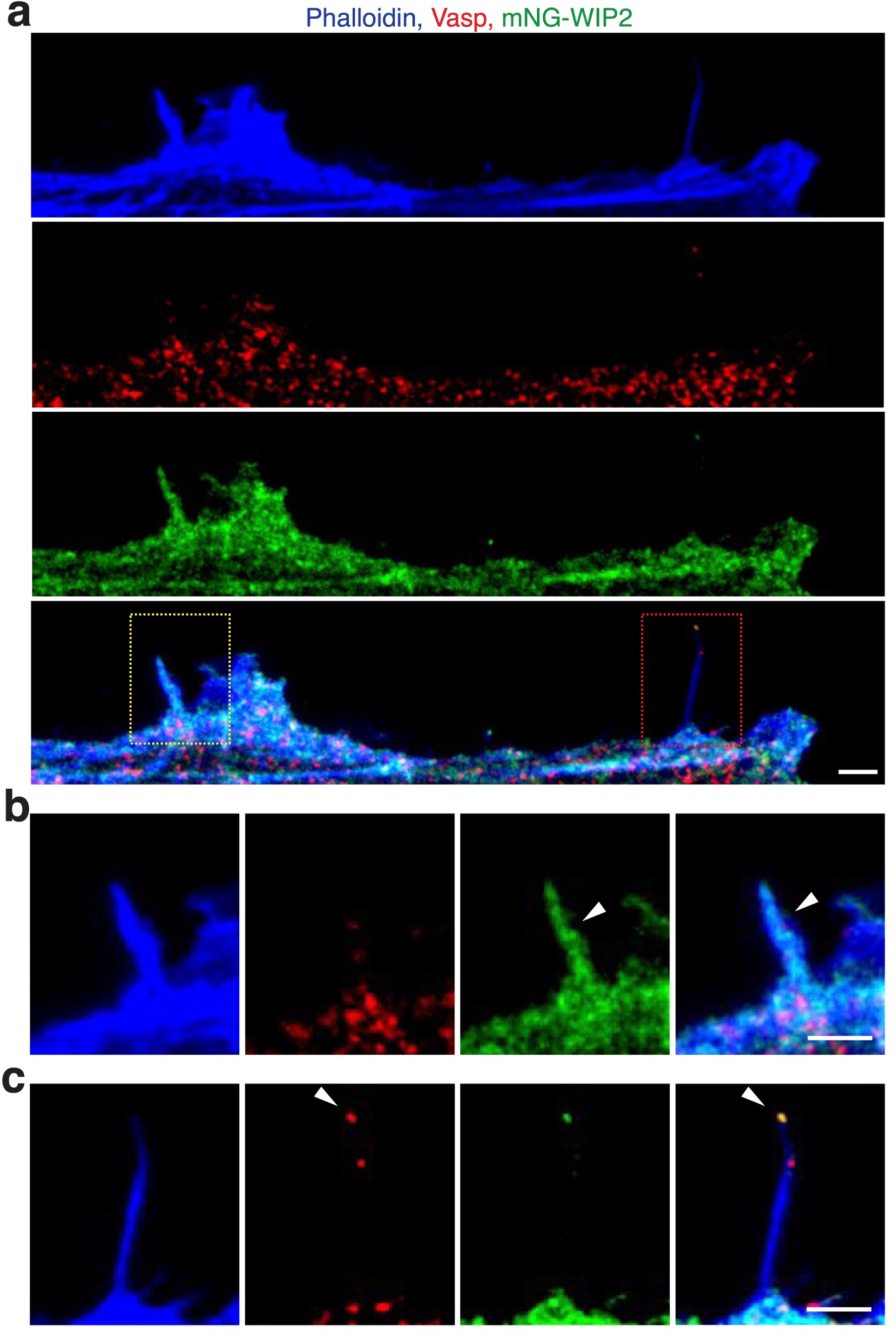
The differentiating C2C12 cells generate both invasive and non-invasive protrusions. **(a)**Immunostaining for Vasp in the differentiating C2C12 cells expressing mNG-WIP2 at 48 hours post MyoD overexpression. The cortical area of a randomly picked cell is shown. **(b)** Enlarged view of yellow-boxed area in **(a)**. Note that the cell projected Vasp^−^ protrusion (invasive) with mNG-WIP2 enriched in the shaft. **(c)** Enlarged view of red-boxed area in **(a)**. Note that the cell projected Vasp^+^ filopodia (non-invasive) without mNG-WIP2 enrichment in the shaft.

**Extended Data Figure 5.**
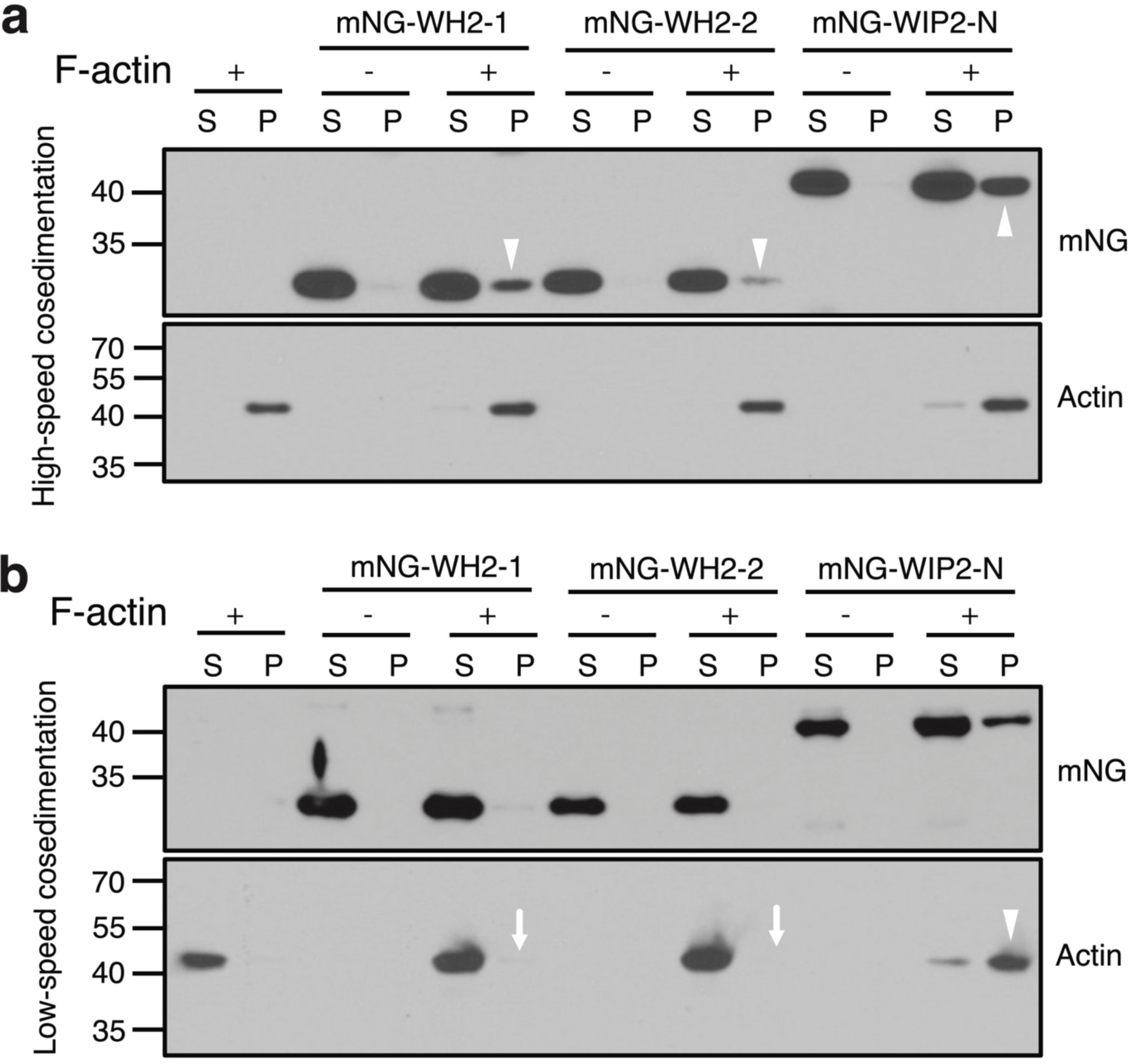
Both WH2 domains of WIP2 are required for actin bundling. **(a-b)** High-speed and low-speed actin co-sedimentation assays for the WH2 domains of WIP2. 4 μM mNG-WH2-1 (mNG tagged first WH2 domain (aa35-53)), mNG-WH2-2 (mNG tagged second WH2 domain (aa80-105)), or mNG-WIP2-N (mNG tagged N-terminal domain of WIP2 containing both WH2-1 and WH2-2 (aa1-176)) was incubated with or without 5 μM F-actin at RT. After 30 minutes, the samples were subjected to high speed (**a**; 100,000g) and low speed (**b**; 13,600g) co-sedimentation assays. After centrifugation, the supernatant (S) and pellet (P) were subjected to SDS-PAGE. In (**a**), all three purified proteins bound F-actin (present in the pellet only when incubated with F-actin) (arrowheads point to the bands in the P lanes), although mNG-WH2-2 showed relatively weaker binding compared to mNG-WH2-1. In (**b**), F-actin was bundled by mNG-WIP2-N (present in the P lane, arrowhead), but not by mNG-WH2-1 or mNG- WH2-2 (absent in the P lane, arrows). Three independent experiments were performed for (**a**) and (**b**) with similar results.

**Extended Data Figure 6.**
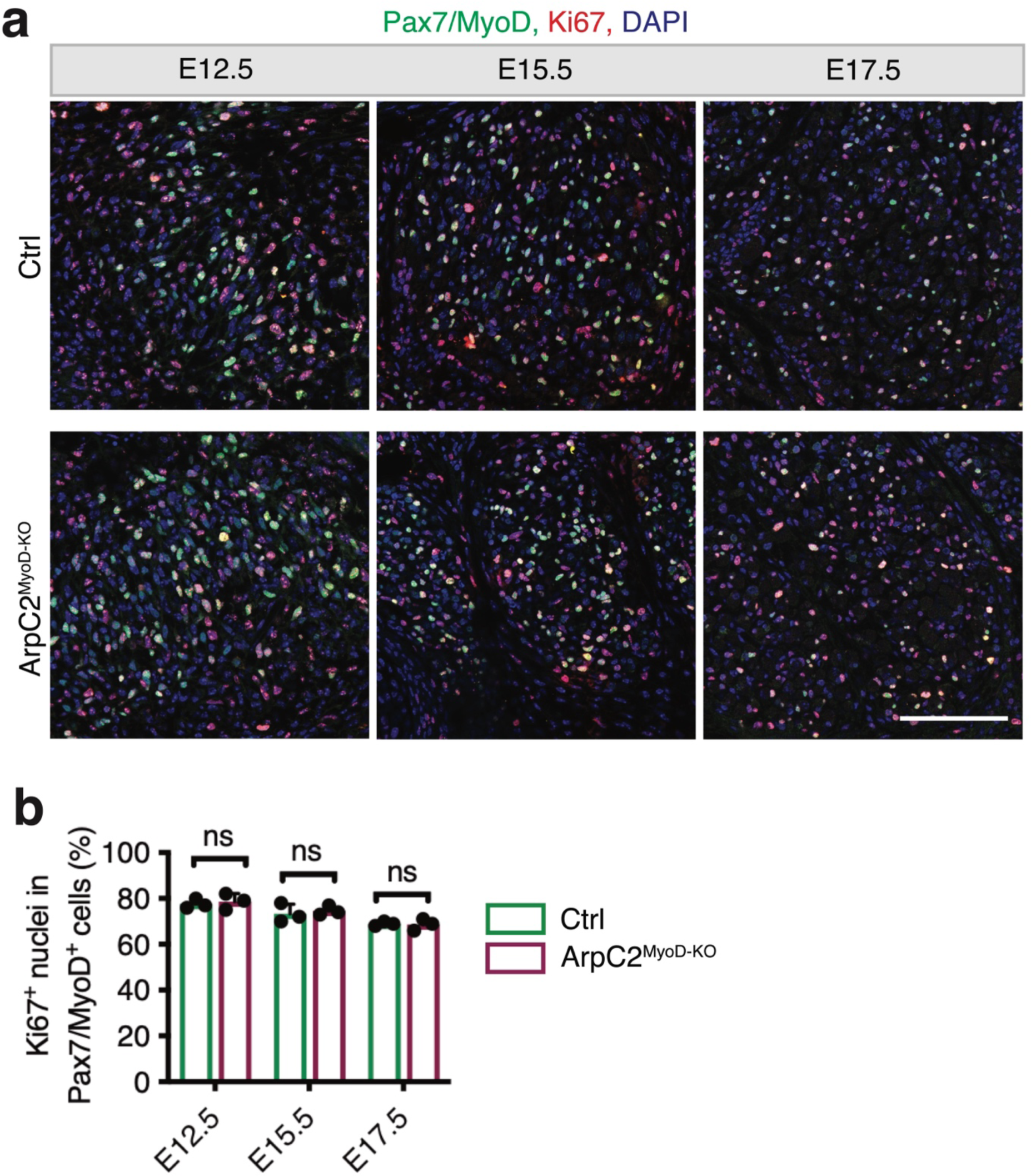
Myoblast proliferation is normal in myoblast specific ArpC2 conditional KO mouse embryos. **(a)**Immunostaining for Ki67 (proliferation maker), MyoD and Pax7 (markers for the myoblasts progenitors) in cross sections of the forelimbs in control and ArpC2^MyoD-KO^ embryos at E12.5, E15.5, and E17.5. Scale bar: 100 μm. **(b)** Quantification of the proliferating myoblasts in the samples shown in (**a**). The percentage of Ki67^+^Pax7/MyoD^+^ nuclei in Pax7/MyoD^+^ nuclei was quantified. Note that Ki67 was normally expressed in the myoblast progenitors of the ArpC2 mutant embryo. Three control and ArpC2^MyoD-KO^ mutant embryos were examined for each time point, respectively. Myoblasts in 12 40x microscopic fields of each embryo were examined. Mean ± s.d. values are shown in the bar graph, and significance was determined by two-tailed student’s t-test. ns: not significant.

**Extended Data Figure 7.**
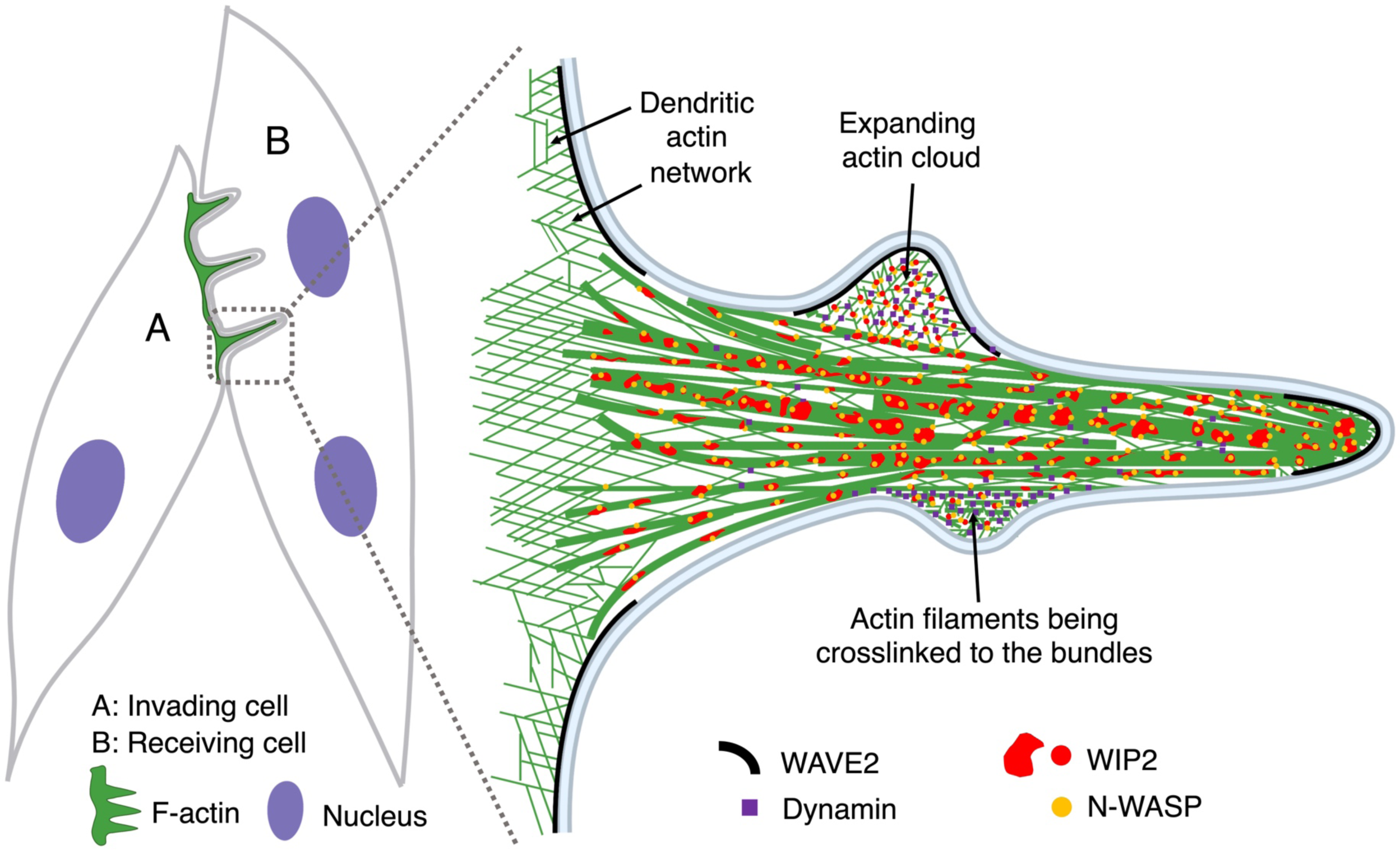
Schematic model on invasive protrusion formation at mammalian fusogenic synapse. Mammalian myoblast fusion occurs at an asymmetric fusogenic synapse, where the invading cell (cell A) generates F-actin-propelled finger-like protrusions to invade the receiving cell (cell B). The individual invasive protrusions arise from the dendritic actin network of the lamellipodia generated by WAVE2, the latter of which is enriched underneath the plasma membrane. N-WASP, on the other hand, is enriched on the F-actin bundle in the shaft via its interaction with WIP2. During protrusion growth, WAVE2 and N-WASP activate the Arp2/3 complex to generate actin clouds at the periphery of the F-actin bundles by polymerizing branched actin filaments. The nascent branched actin filaments are organized into bundles by the large GTPase dynamin. Through the dynamic assembly and disassembly of dynamin in the presence of GTP, all the new actin filaments will be brought into the main F-actin bundles, where WIP2 is enriched and helps to stabilize. Thus, through the coordinated action of two Arp2/3 NPFs (WAVE2 and N-WASP) and two actin-bundling proteins (dynamin and WIP2), the mechanical strength of the branched actin filaments is greatly enhanced, such that the actin bundles can push the plasma membrane forward to generate invasive protrusions.

## Supplementary Video Legends

**Supplementary Video 1. Myoblast fusion occurs between C2C12 cells containing high level of F-actin.**

Time-lapse imaging of C2C12 myoblast fusion at day two in differentiation medium (DM) on a micropattern. SiR-Actin was added to the medium for 30 minutes to label the cellular F-actin prior to live imaging. Scale bar, 100 μm.

**Supplementary Video 2. Invasive membrane protrusions mediate C2C12 cell fusion.**

Time-lapse imaging of fusion between GFP^+^ and GFP^-^ C2C12 cells at day two in DM on a micropattern. Note the appearance of three invasive membrane protrusions (arrowheads) at the fusion site immediately before cell membrane fusion, indicated by GFP transfer from the invading cell to the receiving cell. Scale bar, 10 μm.

**Supplementary Video 3. F-actin is enriched within the invasive membrane protrusions at the fusogenic synapse of C2C12 cells.**

Time-lapse imaging of fusion between an F-tractin-mCherry/GFP co-expressing C2C12 cell and a F-tractin-mCherry expressing C2C12 cell at day two in DM on a micropattern. Note that F-actin was enriched in the invasive protrusion (arrowhead) at the fusion site and dissolved immediately after cell membrane fusion, indicated by GFP transfer (arrow) from the invading cell to the receiving cell. Scale bar, 10 μm.

**Supplementary Video 4. F-actin is enriched at satellite cell fusion sites.**

Time-lapse imaging of a fusion event between two mouse satellite cells co-expressing LifeAct-mScar and FS-EGFP cultured for one day in DM. Note the F-actin enrichment at the invasive front (arrowheads) of the invading cell prior to fusion and its disappearance immediately after fusion. Scale bar, 20 μm.

**Supplementary Video 5. F-actin propelled invasive membrane protrusions mediate the fusion of MyoD-overexpressing C2C12 cells in growth medium.**

Time-lapse imaging of a fusion event between two C2C12 cells co-expressing F-tractin-mCherry and FS-GFP in growth medium (GM) at day two post MyoD overexpression. Note that F-actin was enriched within the finger-like invasive protrusions at the fusogenic synapse (arrows) and dissolved immediately after cell membrane fusion, indicated by the transfer of FS-GFP from the invading cell to the receiving cell. Scale bar, 3 μm.

**Supplementary Video 6. N-WASP, WIP2 and Arp2, but not WAVE2, are enriched at the fusogenic synapse of C2C12 cells.**

Time-lapse imaging of fusion events between C2C12 cells co-expressing LifeAct-mNG and mScar-N-WASP (mScar-WIP2, Arp2-mScar, or mScar-WAVE2) in GM at day two post MyoD overexpression. Note that N-WASP, WIP2 and Arp2, but not WAVE2, enriched with F-actin prior to fusion, and dissolved after fusion was completed. The sites of LifeAct-mNG enrichment are indicated by arrowheads. Scale bar, 5 μm.

**Supplementary Video 7**. **N-WASP is enriched in the shaft of straight protrusions.** Time-lapse imaging of protrusion formation in C2C12 cells co-expressing LifeAct-mNG and mScar-N-WASP in GM at day two post MyoD overexpression. Note that N-WASP was enriched with F-actin along the shaft of straight protrusions. The enrichment of LifeAct-mNG and mScar-N-WASP on randomly picked protrusions are indicated by green and red arrowheads, respectively. Scale bar, 5 μm.

**Supplementary Video 8**. **WIP2 is enriched in the shaft of straight protrusions.** Time-lapse imaging of protrusion formation in C2C12 cells co-expressing LifeAct-mNG and mScar-WIP2 in GM at day two post MyoD overexpression. Note that, similar to N-WASP, WIP2 was enriched with F-actin along the shaft of straight protrusions. The enrichment of LifeAct-mNG and mScar-WIP2 on the protrusions are indicated by green and red arrowheads, respectively. Scale bar, 5 μm.

**Supplementary Video 9. N-WASP/WIP2 and WAVE2 are required to generate invasive protrusions.** Time-lapse imaging of control, N-WASP^-/-^, WIP2^-/-^, WAVE2^-/-^, and Arp2 KD C2C12 cells expressing LifeAct-mNG at day two post MyoD overexpression in GM. Note the straight/stiff vs. bendy/soft finger-like protrusions projected from the leading edge of the lamellipodia in control vs. N-WASP^-/-^ cells; the absence of lamellipodia in WIP2^-/-^, WAVE2^-/-^, and Apr2 KD cells; the bendy/soft protrusions in WIP2^-/-^ cell; the decreased number of protrusions in WAVE2^-/-^ cell; and no protrusion formation in Arp2 KD cells. Scale bar, 10 μm.

**Supplementary Video 10. Pharmacological inhibition of N-WASP impairs the formation of finger-like invasive protrusions.**

Time-lapse imaging of the cortex of a C2C12 cells expressing LifeAct-mNG at day two post MyoD overexpression in GM, immediately after adding DMSO (0.05%) or the N-WASP (5 μM) inhibitor WISK into the medium. Scale bar, 10 μm.

**Supplementary Video 11. WIP2 bundles Arp2/3-mediated branched actin filaments.**

Time-lapse TIRF imaging of actin bundle formation mediated by mNG-WIP2-N, and Shi (*Drosophila* homolog of human dynamin). The TIRF assay was performed in NEM-myosin II-coated flow chambers.

**Supplementary Video 12**. **Visualizing WIP2 bundling actin filaments at the single filament level.**

Time-lapse TIRF imaging of mNG-WIP2-N mediated bundling of actin filaments at the single-filament level. The TIRF assay was performed with NEM-myosin II-coated flow chambers. mNG-WIP2-N was added into the reaction mix of pre-assembled actin filaments. Left panel shows the bundling of two connected F-actin filaments. Middle and right panels show the bundling of two unconnected F-actin filaments. In each panel, the two pre-bundled actin filaments are indicated by green or red arrowheads, and the resulting bundle is indicated by yellow arrowheads. Scale bar, 2 μm.

**Supplementary Video 13. Arp2/3 complex initiates new branched actin polymerization for protrusion initiation and growth.**

Time-lapse imaging of protrusion formation in C2C12 cells co-expressing LifeAct-mNG and Arp2-mScar in GM at day two post MyoD overexpression. Note the Arp2-mScar “budded” out of the sides of an actin bundle (red arrowhead), leading to the formation of an actin cloud. Once a new bundle started to form within the actin cloud (green arrowhead), Arp2 continued to polymerize new branched actin filaments at the periphery of the bundle (red arrowheads), which contributed to the growth of the actin bundle. Scale bar, 3 μm.

**Supplementary Video 14. Dynamin 2 crosslinks new actin filaments into the F-actin bundles during protrusion growth.**

Time-lapse imaging of protrusion growth in C2C12 cells co-expressing LifeAct-mNG and mScar-dynamin 2 in GM at day two post MyoD overexpression. Dynamin 2 was first seen enriched with the actin cloud surrounding the F-actin bundle, until the new actin filaments integrated into the actin bundle, making the bundle thicker and longer. The actin cloud and mScar-dynamin 2 enrichment are indicated by green and red arrowheads respectively. Scale bar, 3 μm.

**Supplementary Video 15. The Arp2/3 complex and Dynamin 2 are co-enriched during protrusion growth.**

Time-lapse imaging of a growing protrusion in C2C12 cells co-expressing Arp2-mNG and mScar-dynamin 2 in GM at day two post MyoD overexpression. Note the co-localization of Arp2 and dynamin 2 during protrusion growth. The enrichment of Arp2-mNG and mScar-dynamin 2 on the protrusion are indicate by green and red arrowheads, respectively. Scale bar, 3 μm.

**Supplementary Video 16. Pharmacological inhibition of dynamin 2 impairs the formation of invasive protrusions.**

Time-lapse imaging of C2C12 cells expressing LifeAct-mNG at day two post MyoD overexpression in GM. 0.05% DMSO or 5 μM dynamin inhibitor MiTMAB (dissolved in DMSO) were added to the medium 24 hours before the live imaging. Note that the MiTMAB caused persistent actin clouds along the protrusions (indicated by arrows), whereas the control cells generated tightly organized actin bundles (indicated by arrowheads).

## Material and Methods

### Mouse Genetics

All animals were maintained under the ethical guidelines of the UT Southwestern Medical Center Animal Care and Use Committee according to NIH guidelines. C57BL/6J were purchased from The Jackson Laboratory (000664). The MyoD^iCre^ and ArpC2^flox/flox^ mice were generously provided by Drs. David Goldhamer and Rong Li, respectively. For conditional knockout of ArpC2 in embryonic myogenic progenitors, MyoD^iCre^ mice were crossed with ArpC2^flox/flox^ mice to generate MyoD^iCre^; ArpC2^flox/wt^ and ArpC2^flox/wt^ progenies, which were further crossed to generate the control (MyoD^iCre^; ArpC2^wt/wt^) and mutant (MyoD^iCre^; ArpC2^flox/flox^) genotypes used in this study. The control and mutant littermates were used in each cohort of experiment. The mouse embryos were staged according to Kaufman^63^. Noon on the day when a vaginal plug was observed was taken as embryonic day 0.5 of gestation (E0.5).

### Satellite Cell Isolation and Cell Culture

Satellite cells were isolated from limb skeletal muscles of wild-type C57BL/6J male mice at the age of eight weeks. Briefly, muscles were minced and digested in 800U/ml type II collagenase (Worthington; LS004196) in F-10 Ham’s medium (ThermoFisher Scientific; 11550043) containing 10% horse serum at 37 °C for 90 minutes with rocking to dissociate muscle fibers and dissolve connective tissues. The dissociated myofiber fragments were collected by centrifuge and digested in 0.5U/ml dispase (Gibco; 17105041) in F-10 Ham’s medium at 37 °C for 30 minutes with rocking. Digestion was stopped with F-10 Ham’s medium containing 20% FBS. Cells were then filtered from debris, centrifuged and resuspended in satellite cell growth medium (SCGM: F-10 Ham’s medium supplemented with 20% FBS, 4 ng/ ml FGF2, 1% penicillin–streptomycin and 10mM HEPEs). The cell suspension from each animal was pre-plated twice into the regular 100 mm tissue culture-treated dishes for 30 minutes at 37 °C to eliminate fibroblasts. The supernatant containing mostly myoblasts was then transferred into collagen-coated dishes for culture in SCGM. C2C12 cells and U2OS cells were maintained in growth medium (GM; DMEM supplemented with 10% FBS, 1% penicillin/streptomycin and 10mM HEPEs). For low serum induced myogenic differentiation, C2C12 cells and satellite cells were cultured in differentiation medium (DM; DMEM supplemented with 2% horse serum, 1% penicillin-streptomycin and 10mM HEPEs). Cells in all the conditions were cultured at 37 °C with 5% CO2.

### MyoD-induced Myoblast Fusion in Growth Medium

For live cell imaging of the MyoD^OE^ cells, C2C12 cells were seeded on fibronectin (Sigma; F1141) coated glass coverslips at 30% confluency in GM. After one day, retrovirus containing MyoD was mixed with polybrene (7 μg/ml) and added to GM to infect the cells for a day. Subsequently, the infected cells were incubated in fresh GM for another day and subjected to live cell imaging. To analyze fusion index of the MyoD^OE^ cells, C2C12 cells were seeded in tissue culture-treated 6-well plates at 60% confluency in GM. After one day, the cells were infected by retrovirus containing MyoD for a day, followed by a fresh GM change. After another day, the cells were harvested for immunostaining and/or western blot.

### Pharmacological Treatments of C2C12 cells

To pharmacologically inhibit actin polymerization during myoblast fusion, the 100% confluent C2C2 cells were differentiated in DM for 48 hours, followed by incubation with DMSO (0.05%), cytochalasin D (18nM), Arp2/3 inhibitor (50 μM) or formin inhibitor SMIFH2 (10 μM) for 24 hours. Then, the cells were fixed in 4% paraformaldehyde (PFA) and immunostained with anti-MyoG and anti-MHC to assess their differentiation and fusion index.

To observe the immediate effect of N-WASP inhibitor, WISK, on the cortical protrusion formation, C2C12 cells expressing LifeAct-mNG were cultured on fibronectin coated glass coverslips at 30% confluency. After 24 hours, the cells were infected by retrovirus containing MyoD. 48 hours post MyoD overexpression, the cells were subject to live cell imaging in GM containing DMSO (0.05%) for one hour. Subsequently, the medium was quickly replaced with fresh GM with WISK (5 μM) and the cells were immediately subjected to live imaging for one hour.

To observe the effect of MiTMAB on the cortical protrusion formation, LifeAct-mNG expressing C2C12 cells were cultured on fibronectin coated glass coverslips at 30% confluency. After 24 hours, the cells were infected by retrovirus containing MyoD. 24 hours post MyoD overexpression, the cells were cultured in GM containing DMSO (0.05%) or MiTMAB (5 μM). After another 24 hours, the treated cells were subjected to live imaging.

### Immunocytochemistry

Cultured cells were fixed in 4% PFA for 15 minutes at room temperature (RT). After extensive washes with PBS, the cells were incubated with blocking buffer (PBS containing 2% BSA and 0.1% TritonX-100) for 20 minutes at RT. Then the cells were incubated with primary antibodies diluted in the blocking buffer at 4°C overnight. After washes with PBS, the cells were further incubated with the appropriate alexa fluor-conjugated secondary antibodies diluted in the blocking buffer for one hour at RT. The cells were then washed with PBS and imaged using a Leica TCS SP8 inverted microscope. The following primary antibodies were used: rabbit anti-WAVE2 (1:200; Cell Signaling Technologies; 3659), mouse anti-MyoG (1:30; DSHB; F5D), mouse anti-MHC (1:100; DSHB; MF20), mouse anti-dynamin 2 (Santa Cruz; sc-166669) and mouse anti-Vasp (1:100, Santa Cruz; sc-46668). The alexa fluor 488-, 568- and 647- conjugated (Invitrogen) secondary antibodies were used at 1:200. For F-actin labelling, alexa fluor 405-, 488-, 568- or 647-conjugated phalloidin (Invitrogen) was used at 1:200 for one hour at RT.

### Immunohistochemistry

Mouse embryos were fixed in 4% PFA at 4°C for three days. Whole mount forelimb staining was performed using alkaline phosphatase (AP) conjugated myosin antibody as described in a previous study^64^. Briefly, the skinned forelimbs were post-fixed in 100% methanol prior to the staining. After extensive washes with PBST (PBS containing 0.1% Tween-20), the limbs were incubated in PBST for one hour at 70°C to inactivate endogenous AP, followed by bleaching with 6% Hydrogen Peroxide in PBST for one hour at RT. The bleached samples were then washed in PBST and incubated in the blocking solution (0.1% Triton; 1% BSA; 0.15% glycine in PBS) at RT with rocking. The limbs were subsequently incubated with anti-myosin-AP (My32) (1:800, Sigma; A4335) in blocking solution overnight at 4°C with rocking, followed by extensive washes with PBST. Next, the limbs were washed in NTMT (100mM NaCl, 100mM Tris-HCl (pH 9.5), 50mM MgCl2, 0.1% Tween-20) for 15 minutes and developed with NBT/BCIP substrate solution (ThermoFisher Scientific; 34042) at RT. For immunofluorescent staining of muscle tissue sections, the PFA fixed embryos were dehydrated in 30% sucrose at 4°C overnight. The specimens were embedded in Tissue-Plus O.C.T. Compound (Fisher Scientific; 23-730-571) and 12-μm cryosections were collected onto Superfrost Plus Microscope Slides (Fisher Scientific;12-550-15). Then, the cryosections were incubated with blocking buffer for 20 minutes at RT, followed by overnight incubation with mouse anti-MyoG (1:30; DSHB; F5D) and/or mouse anti-MHC (1:100; DSHB; MF20) at 4°C overnight. After extensive washing with PBS, the sections were incubated with alexa fluor-conjugated antibodies for one hour at RT. Subsequently, the sections were washed with PBS and subjected to imaging using a Leica TCS SP8 inverted microscope.

### CRISPR sgRNA Knockout in Myoblasts

The sgRNAs targeting the mouse *N-WASP*, *WIP2* and *WAVE2* open reading frames were designed using the online software CRISPOR. The sgRNAs were individually cloned into lenti-CRISPR v2 vector (Addgene; 52961). The sgRNA sequences are as follows:

N-WASP gRNA, 5’ CCGCGGAGGGTCACCAACGT 3’;

WAVE2 gRNA, 5’ TCCAGCTCGCTTGTATCGCT 3’;

WIP2 gRNA, 5’ AGCACTCCGATCATTAACGT 3’.

The lenti-CRISPR v2 constructs containing specific sgRNAs were transfected into Lenti-X 293T cells (Takara; 632180) with psPAX2 and VSV-G plasmids using the FuGENE HD transfection reagent (Promega; E2311) to produce lentivirus. Two days after transfection, the lentivirus supernatants were filtered and mixed with polybrene (7 μg/ml) to infect the C2C12 cells. Seven days post infection, single cell clones were isolated and expanded. Western blot was used to detect the expression level of target proteins to select the knockout cell clones.

### Retroviral Vector Preparations and Expression

The full-length open reading frames of mouse *MyoD*, *MyoG*, *N-WASP*, *WAVE2*, *WIP2* and *Arp2* were amplified using cDNAs generated from C2C12 cells at day two in DM. LifeAct, mNeongreen, mScarleti, dynamin 2, and F-tractin-mCherry were amplified from LifeAct-Neongreen (Addgene; 98877), LCK-mScarleti (Addgene; 98821), Dyn2-pmCherryN1 (Addgene; 27689) and C1-F-tractin-mCherry (Addgene; 155218), respectively. All the constructs were assembled into the retroviral vector pMXs-Puro (Cell Biolabs; RTV-012) using the NEBuilder HiFi DNA Assembly Cloning Kit (NEB; E2621L). The retroviral construct containing farnesylation signal-GFP was purchased from addgene (21836). MyoD and MyoG were used to induce C2C12 cell fusion in GM. LifeAct-mScarleti, farnesylation signal-GFP, and F-tractin-mCherry were used to observe the invasive finger-like protrusions during C2C12 cell and satellite cell fusion. Arp2-mNeongreen, Arp2-mScarleti, mNeongreen-N-WASP, mScarleti-N-WASP, mScarleti-WAVE2, mNeongreen-WIP2, mScarleti-WIP2, and mScarleti-dynamin2 were constructed to visualize the subcellular localization of the corresponding tagged proteins. To package the retrovirus, two micrograms of retroviral plasmid DNA was transfected into platinum-E cells (Cell Biolabs; RV-101) using the FuGENE HD transfection reagent. Two days after transfection, the virus-containing medium was filtered, mixed with polybrene (7 μg/ml), and used to infect cells. One day after infection, cells were washed with PBS and cultured in GM. For rescue experiments, sgRNA-insensitive DNA cassettes for the target genes were used.

### shRNA-based Gene Silence

ShRNAs against *N-WASP*, *WIP2*, *WAVE2* and *Arp2* were individually cloned to lentiviral RNAi construct LENC (Addgene; 111163) or Tet-On lentiviral RNAi construct RT3REN (Addgene; 111166)^65^. The shRNA sequences used in this study are as follows:

Sh1N-WASP: TAGTCACACATTTCTTGCCGAG

Sh2N-WASP: TTTTGCTTCTTCTTCATTGGCA

Sh1WIP2: TAGTGAACAATTAAAAGTCCGG

Sh2WIP2: TTAGTGTGTATGCACTGCTCAT

Sh1WAVE2: TTATCTTTCTTTTCTTTCCTAT

Sh2WAVE2: TAACATTATTTGAAGTAGCTAA

ShArp2: TTTCTTGGTACTCTTGTCTGGT

The constructs were co-transfected with psPAX2 and VSV-G plasmids into Lenti-X 293T cells using the FuGENE HD transfection reagent to produce lentivirus. Two days after transfection, the medium containing lentiviruses were harvested, mixed with polybrene (7 μg/ml) and used to infect the C2C12 cells. Two days later, puromycin (2 μg/ml) was added to GM for three days to select the infected cells. The surviving cells were subsequently expanded in GM and seeded at 60% confluence in tissue culture-treated 6-well plates. For straight knockdown experiments, when cells grew to 100% confluence, the GM was switched to DM. The cells were then allowed to differentiate for five days during which their differentiation and fusion were assessed. For doxycycline-induced shRNA expression, doxycycline (1 μg/ml) or empty vehicle was added to GM 24 hours after the cells were seeded. 48 hours post doxycycline addition, the GM was switched to DM with doxycycline (1 μg/ml) or empty vehicle. The cells were then allowed to differentiate for five days during which their differentiation and fusion were assessed.

### Western Blot

For western blot, cells were lysed by ice-cold RIPA buffer (150mM NaCl, 1% NP40, 0.1% SDS and 50mM Tris, PH7.4) containing protease and phosphatase inhibitor (Cell Signaling Technologies; 5872) for 20 minutes. The supernatants were collected by centrifugation at 140,000 x g for 15 minutes. Protein concentrations were determined using the Bradford Protein Assay Kit (Bio-Rad; 5000201). 10-30μg total protein was loaded and separated by 10% SDS-PAGE gel and transferred to PVDF membranes (Millipore; GVHP29325). Then, the membranes were blocked for one hour at RT in PBS containing 5% nonfat dry milk and 0.1% Tween-20 (PBSBT) and subsequently were incubated with primary antibodies diluted 1:1000 in PBSBT overnight at 4°C. The membranes were then washed with PBST and incubated with appropriate HRP-conjugated secondary antibodies diluted in PBSBT for one hour at RT. After extensive washes with PBST, the membranes were developed with the ECL western blotting substrate (ThermoFisher Scientific; 32209). The following primary antibodies were used: mouse anti-WASP (1:1000; Santa Cruz; 4860), rabbit anti-N-WASP (1:1000; Cell Signaling Technologies; 4848), mouse anti-WIP1 (1:1000, Santa Cruz; sc-271113), rabbit anti-WIP2 (1:1000; ThermoFisher Scientific; PA5-55086), mouse anti-WAVE1 (1:1000; Santa Cruz; sc-271507), rabbit anti-WAVE2 (1:1000; Cell Signaling Technologies; 3659), rabbit anti-WAVE3 (1:1000; Cell Signaling Technologies; 2806), mouse anti-Arp2 (1:1000; Santa Cruz; sc-166103), mouse anti-Arp3 (1:1000; Santa Cruz; sc-48344), mouse anti-MyoG (1:30; DSHB; F5D), mouse anti-MHC (1:200; DSHB; MF20), rabbit anti-ABI1 (1:1000; Cell Signaling Technologies; 39444), rabbit anti-Cyfip1(1:1000; Cell Signaling Technologies; 81221), mouse anti-mNeongreen (Proteintech; 32F6), mouse anti-actin (Sigma: A4700), sheep anti-ESGP (1:1000; R&D Systems; AF4580) and rabbit anti-β-Tubulin (1:1000; Cell Signaling Technologies; 2146).

### Protein Purification

Three partial sequences of mouse WIP2, aa1-176 (WIP2-N), aa35-53(WH2-1) and aa80-105 (WH2-2), were fused with mNeongreen at their C-terminus with a flexible peptide linker and cloned into pET28a vector (Millipore; 69864). Protein expression in *Escherichia coli* BL21(DE3) cells was induced by 200 μM IPTG (OD600=0.8) for 18 hours at 16°C. Bacteria were collected and lysed by sonication in lysis buffer (50 mM Tris-HCl pH 8.0, 500 mM NaCl, 5 mM imidazole, 5 mM β-mercaptoethanol) supplemented with protease inhibitors, followed by centrifugation at 20,000 rpm for 45 minutes at 4°C. The proteins were then purified with Ni-NTA agarose beads (QIAGEN; 30210) according to the manufacturer’s instructions. Proteins eluted by 100 mM imidazole were collected and then desalted with disposable PD 10 desalting columns (Sigma, GE17-0851-01) against stocking buffer (20 mM HEPES (pH 7.3), 150 mM KCl, 1 mM EGTA and 1 mM dithiothreitol). *Drosophila* dynamin was purified as described in our previous study^21^. The purified proteins were subjected to SDS-PAGE gels and coomassie blue staining to determine the purity of the proteins. Bradford assays were performed to determine the protein concentrations. The purified proteins were then flash frozen in liquid nitrogen and stored at −80 °C until use.

### F-actin Co-sedimentation Assay

F-actin co-sedimentation assays were performed to determine the F-actin binding and bundling activities of mouse WIP2. F-actin pre-assembled from 5 μM G-actin was incubated alone or with the indicated concentrations of proteins in 5 mM HEPES (pH 7.3), 50 mM KCl, 2 mM MgCl2, 0.5 mM EGTA, 3 mM imidazole, 60 nM ATP, 60 nM CaCl2, 0.4 mM DTT and 0.003% NaN3 for 30 minutes at RT. The samples were then spun for 30 minutes at 4 °C at 100,000g (high-speed co-sedimentation) or 13,600g (low-speed co-sedimentation). After centrifugation the supernatants and pellets were resolved by 10% SDS-PAGE and the gels were stained with coomassie blue or subjected to western blot.

### Pharmacological Treatments of Dynamin-actin Interaction

To determine the effect of dynamin inhibitor, MiTMAB, on dynamin-actin interaction *in vitro*, F-actin pre-assembled from 1 μM G-actin was incubated alone or with 1μM His-Shibirie in 5 mM HEPES (pH 7.3), 50 mM KCl, 2 mM MgCl2, 0.5 mM EGTA, 3 mM imidazole, 60 nM ATP, 60 nM CaCl2, 0.4 mM DTT and 0.003% NaN3 for 30 minutes at RT. Subsequently, 2mM GTP with or without 40 μM MiTMAB was added to the appropriate reaction mixtures and incubated for five minutes at RT. Then, the samples were mixed with alexa fluor 488-conjugated phalloidin (Invitrogen), dropped on the glass coverslips, and subjected to imaging using a Leica TCS SP8 inverted microscope.

### Time-lapse Imaging and Analysis

Time-lapse imaging was performed with a Nikon A1R confocal microscope using a 40× (0.4 NA) objective with 37 °C and 5% CO2 using a Nikon Biostation CT. To image C2C12 fusion in DM, cells were seeded on micropatterns (with a diameter of 250 μm) and imaged at 48 hours after switching from GM to DM. The micropatterns were generated as described previously ^66^. To image the fusion of mouse satellite cells in DM, the cells seeded on the fibronectin-coated cover glass (MATTEK; P35G-0-14-C) were imaged at 24 hours after switching from GM to DM. To image MyoD^OE^-induced myoblast fusion in GM, C2C12 cells were seeded on the fibronectin-coated cover glass and imaged at 48 hours post MyoD overexpression. The cells were imaged at one-to two-minute interval. After time-lapse imaging, ImageJ (NIH, 64-bit Java 1.8.0_172) was used to project the z-stacks in 2D, using maximum intensity projection and the resulting 2D images were assembled into a time-lapse video.

### Total Internal Reflection Fluorescence (TIRF) Microscopy

Flow chambers were assembled and coated with 10 nM NEM-myosin II in high salt TBS buffer (50 mM Tris-HCl (pH 7.5) and 600 mM NaCl) for one minute, washed with high salt BSA (1% BSA, 50 mM Tris-HCl (pH 7.5) and 600 mM NaCl) and low salt BSA (1% BSA, 50 mM Tris-HCl (pH 7.5) and 150 mM NaCl) sequentially, then washed with the TIRF buffer (50 mM KCl, 1 mM MgCl2, 1 mM EGTA, 10 mM imidazole, 100 mM DTT, 0.2 mM ATP, 15 mM glucose, 20 μg/ml catalase, 100 μg/ml glucose oxidase, 0.2% BSA and 0.5% methylcellulose (pH 7.0)). To visualize WIP2 and dynamin-mediated bundling of the Arp2/3-nucleated branched actin network, 50 nM Arp2/3 (Cytoskeleton; RP01P), 1 μM VCA domain of WASP (Cytoskeleton; VCG03-A), 2 μM G-actin (70% unlabeled and 30% rhodamine-G-actin; cytoskeleton) were mixed in TIRF buffer, loaded into the flow chambers and subjected to Ring-TIRF imaging immediately using a GE OMX-SR super-resolution microscope equipped with a 60X/1.49 UPlanApo oil objective lens. After 20 minutes, 4 μM mNeongreen-WIP2-N or His-Shibire was loaded into the flow chambers while the actin filaments were continuously imaged by Ring-TIRF.

### Super-Resolution Microscopy

To visualize the distribution of WIP2 along actin bundles, 1 μM mNeongreen-WIP-N was incubated with 1 μM F-actin and 1 μM alexa fluor 568-phalloidin in 5 mM HEPES (pH 7.3), 50 mM KCl, 2 mM MgCl2, 0.5 mM EGTA, 3 mM imidazole, 60 nM ATP, 60 nM CaCl2, 0.4 mM DTT and 0.003% NaN3 for 30 minutes at RT. Subsequently, the samples were diluted 20-fold in the fluorescence buffer (0.5% methycellulose, 50 mM KCl, 1 mM MgCl2, 1 mM EGTA, 10 mM imidazole, 100 mM DTT, 0.02 mg/ml catalase, 0.1 mg/ml glucose oxidase, 15 mM glucose, 0.1 mM CaCl2, 0.1 mM ATP and 0.005% NaN3). Then, 4 μl of diluted samples were mounted onto glass slide, covered with a poly-L-lysine-coated coverslip, and subjected to structured illumination microscopy (SIM) using a GE OMX-SR super-resolution microscope equipped with a 60X/1.42 UPlanApo oil objective lens.

### Electron Microscopy

To observe the invasive protrusions at the contact sites of C2C12 cells in DM, C2C12 cells were first cultured on tissue-culture treated 35mm dishes in GM. Once the cells reached 100% confluence, GM was replaced with DM. After 48 hours, the cells were fixed in a solution containing 2.5% glutaraldehyde, 1% sucrose and 3mM CaCl2 in 0.1M sodium cacodylate buffer (pH 7.4) at 4°C overnight. On the following day, samples were rinsed in 0.1M cacodylate buffer containing 3% sucrose and 3mM CaCl2 before being post-fixed for 1.5 hours on ice with 1% osmium tetroxide and 0.8% potassium ferricyanide. En bloc staining was performed by treating the samples with 0.25% tannic acid and 4% uranyl acetate. Then, the samples were dehydrated, infiltrated, and embedded in EPON resin. Using a LEICA ultramicrotome (UC6), resin-embedded samples were cut into 70 nm sections and collected on copper slot grids. These sections were viewed with a JEOL 1400 transmission electron microscope after being post-stained with 2% uranyl acetate and Reynold’s stain. To observe the invasive protrusions at the contact sites of MyoD^OE^ C2C12 cells, C2C12 cells cultured on tissue-culture treated 35mm dishes were infected with retrovirus carrying MyoD in GM. 48 hours post infection, the cells were subjected to fixation, staining and electron microscopy as described above.

To observe the invasive protrusions at the contact sites of myoblasts *in vivo*, hindlimbs of E15.5 mouse embryos were fixed in a solution containing 3% paraformaldehyde, 2% glutaraldehyde, 1% sucrose, 3mM CaCl2 in 0.1M sodium cacodylate buffer (pH 7.4) at 4°C overnight. Samples were subsequently washed with 0.1M cacodylate buffer containing 3% sucrose and 3mM CaCl2, and post fixed with 1% osmium tetroxide in 0.1M sodium cacodylate buffer for 1.5 hours on ice. The limbs were stained with 2% uranyl acetate, dehydrated and embedded in EPON resin. The embedded samples were then cut into 70 nm thick sections using LEICA ultramicrotome (UC6) and collected on copper slot grids. These sections were post-stained with 2% uranyl acetate and sato’s lead solution, and examined using a JEOL 1400 transmission electron microscope.

### Negative-Stain Electron Microscopy

Negative-stain electron microscopy was performed to visualize the distribution of WIP2 and dynamin along F-actin bundles. A total of 5 μM F-actin was incubated with an appropriate concentration of mNeogreen-WIP2 or His-dynamin in 6.5 mM HEPES (pH 7.3), 50 mM KCl, 2 mM MgCl2, 0.3 mM EGTA, 20 nM ATP, 20 nM CaCl2, 0.3 mM DTT and 0.001% NaN3 at RT for 30 minutes. The reaction mixture was loaded onto carbon-coated, glow-discharged 400 mesh copper grids and incubated for five minutes, followed by two quick washes in 1X actin polymerization buffer (50 mM KCl, 2 mM MgCl2, 5 mM guanidine carbonate (pH 7.5), and 1 mM ATP). The grid was stained in two drops of 0.75% uranyl formate for 20 seconds and air dried. Electron micrographs were collected on a JEOL 1400 transmission electron microscope.

### Statistics and Reproducibility

Statistical significance was assessed using two-tailed student’s t-test. *P* values were obtained using GraphPad Prism. The numbers of biological replicates for each experiment are indicated in the figure legends.

## Acknowledgements

We thank the UT Southwestern Animal Resource Center for assistance with the mouse colony maintenance and the UT Southwestern Quantitative Light Microscopy Core Facility for TIRF and SIM microscopy assistance. This work was supported by NIH grants (R01AR075005, R01AR053173, and R35GM136316) to E.H.C., and a Fellowship from the UT Southwestern Hamon Center for Regenerative Science and Medicine (CRSM) to K.H.L.

## Author contributions

Y.L., T.W., B.R. and E.H.C. designed the project. Y.L., T.W., B.R., P.P. and B.L. performed experiments. B.R. and E.H.C collaborated with K.H.L. and D.W.S. on preparing micropatterns for live cell imaging experiments. D.J.G. provided the MyoD^iCre^ mouse line. R.L. provided ArpC2^flox/flox^ mouse line. B.L. and D.P. purified WH2-1, WH2-2 and WIP2-N. R.Z. purified dynamin. Y.L., T.W., B.R. and E.H.C. analyzed the data. Y.L. and E.H.C. made the figures and wrote the manuscript. All authors commented on the manuscript.

## Data availability

The main data supporting the findings of this study are available within the article and its Supplementary Information files. All other data supporting the findings of this study are available from the corresponding author upon reasonable request.

## Author information

The authors declare no competing financial interests. Correspondence and requests for materials should be addressed to E.H.C.

